# Glycerol Metabolism Contributes to Competition by Oral Streptococci through Production of Hydrogen Peroxide

**DOI:** 10.1101/2024.06.28.598274

**Authors:** Zachary A. Taylor, Ping Chen, Payam Noeparvar, Danniel N. Pham, Alejandro R. Walker, Todd Kitten, Lin Zeng

## Abstract

As a biological byproduct from both humans and microbes, glycerol’s contribution to microbial homeostasis in the oral cavity remains understudied. Here we examined glycerol metabolism by *Streptococcus sanguinis,* a commensal associated with oral health. Genetic mutants of glucose-PTS enzyme II (*manL*), glycerol metabolism (*glp* and *dha* pathways), and transcriptional regulators were characterized with regard to glycerol catabolism, growth, production of hydrogen peroxide (H_2_O_2_), transcription, and competition with *Streptococcus mutans*. Biochemical assays identified the *glp* pathway as a novel source of H_2_O_2_ production by *S. sanguinis* that is independent of pyruvate oxidase (SpxB). Genetic analysis indicated that the *glp* pathway requires glycerol and a transcriptional regulator, GlpR, for expression and is negatively regulated by PTS, but not the catabolite control protein, CcpA. Conversely, deletion of either *manL* or *ccpA* increased expression of *spxB* and a second, H_2_O_2_-non-producing glycerol metabolic pathway (*dha*), indicative of a mode of regulation consistent with conventional carbon catabolite repression (CCR). In a plate-based antagonism assay and competition assays performed with planktonic and biofilm-grown cells, glycerol greatly benefited the competitive fitness of *S. sanguinis* against *S. mutans.* The *glp* pathway appears to be conserved in several commensal streptococci and actively expressed in caries-free plaque samples. Our study suggests that glycerol metabolism plays a more significant role in the ecology of the oral cavity than previously understood. Commensal streptococci, though not able to use glycerol as a sole carbohydrate for growth, benefit from catabolism of glycerol through production of both ATP and H_2_O_2_.

**Importance:** Glycerol is an abundant carbohydrate found in oral cavity, both due to biological activities of humans and microbes, and as a common ingredient of foods and health care products. However, very little is understood regarding the metabolism of glycerol by some of the most abundant oral bacteria, commensal streptococci. This was in part because most streptococci cannot grow on glycerol as the sole carbon source. Here we show that *Streptococcus sanguinis*, an oral commensal associated with dental health, can degrade glycerol for persistence and competition through two independent pathways, one of which generates hydrogen peroxide at levels capable of inhibiting a dental pathobiont, *Streptococcus mutans*. Preliminary studies suggest that several other commensal streptococci are also able to catabolize glycerol, and glycerol-related genes are being actively expressed in human dental plaque samples. Our findings reveal the potential of glycerol to significantly impact microbial homeostasis which warrants further exploration.

## Introduction

Glycerol is a byproduct of microbial fermentation of carbohydrates, including what is produced by yeasts as a compatible solute to boost osmo-tolerance, and by certain oral commensal bacteria including *Corynebacterium* (1, 2). Glycerol is widely used as a food additive and important industrial ingredient for many health-care products and cosmetics. In addition, glycerol and related compounds can be released from degradation of dietary or biological lipids, by the action of lipases or phospholipases, and are found in human blood as metabolic intermediates. As such, glycerol is an abundant carbon source for the oral microbiota and can potentially be utilized for both energy production and as a precursor to many essential biomolecules such as phospholipids and lipoteichoic acids that are essential for bacterial envelope biogenesis (3, 4). An increasing body of research has indicated the significant contribution of glycerol metabolism to bacterial fitness and pathophysiology (3, 5–8). For example, glycerol oxidation by the bacterial genus *Mycoplasma* is considered critical to the virulence of these bacteria, in large part due to the resulting production of hydrogen peroxide (H_2_O_2_) (5).

Much of our understanding of glycerol metabolism by Gram-positive bacteria has been derived from research in the model organism *Bacillus subtilis* (9, 10), and in *Enterococcus faecalis*, a gut commensal and an opportunistic human pathogen (11)*. E. faecalis* can utilize glycerol for growth, producing end products such as lactic acid, acetic acid, and ethanol (7). Regarding glycerol fermentation for acid production, a significant proportion of lactic acid bacteria (LAB) have been suggested to lack such capability and there exists notable intra-species variations (12, 13). Two pathways are genetically encoded by enterococci and related LAB for the metabolism of glycerol, dehydrogenation and phosphorylation, often with varying degrees of genomic reduction in glycerol-negative LAB (4). For the dehydrogenation pathway (encoded by *gldA-dhaKLM*), glycerol is converted into dihydroxyacetone (DHA) by a glycerol dehydrogenase (*gldA*) with a concomitant reduction of NAD^+^ to NADH, and then into dihydroxyacetone phosphate (DHAP) by a DHA kinase (*dhaKLM*), which depends on bacterial PTS (phosphoenolpyruvate-dependent sugar: phosphotransferase system)(14) for supply of the phosphoryl group. For the phosphorylation pathway (*glpKOF*), glycerol is first phosphorylated by a glycerol kinase (GlpK; EC 2.7.1.30) into glycerol-3-phosphate (Gly-3-P), then oxidized, in the presence of oxygen, into DHAP and H_2_O_2_ by the gene product of *glpO* (Gly-3-P oxidase; EC 1.1.3.21). Genes coding for both metabolic pathways in *E. faecalis* are important for glycerol metabolism and are expressed in a strain-dependent manner (13), and both pathways were required for virulence of *E. faecalis* in a mouse intraperitoneal model (8). The glycerol dehydrogenation pathway is considered by some to be indispensable for efficient glycerol metabolism in *E. faecalis* as glycerol phosphorylation creates H_2_O_2_ as a byproduct, accumulation of which could prove detrimental to the very bacterium that produces it (4). Conversely, it has been shown that oxygen is necessary for acid production from glycerol by most LAB (12, 15), which differs from the metabolism of other common carbohydrates such as glucose, and that genes for the phosphorylation pathway appear to be better conserved overall in LAB than those of the dehydrogenation pathway (4). Furthermore, several species of the viridans streptococci that are considered important to oral health, including *Streptococcus mutans, Streptococcus gordonii, Streptococcus sanguinis, Streptococcus mitis,* and *Streptococcus oralis*, cannot ferment glycerol to lower the pH or support bacterial growth (12, 16). Nonetheless, a recent study has suggested that *S. sanguinis* can apparently utilize glycerol to achieve certain morphological phenotypes (17, 18). While some of these streptococcal species, including the major etiologic agent of dental caries, *S. mutans,* apparently lack critical genes required for either glycerol pathway, the *S. sanguinis* SK36 genome harbors the complete sets of genes for both (Fig. 1). The function and significance of these conserved genetic pathways in *S. sanguinis*, namely *glpKOF* and *gldA-dhaKLM,* remain to be determined.

**Fig. 1.**
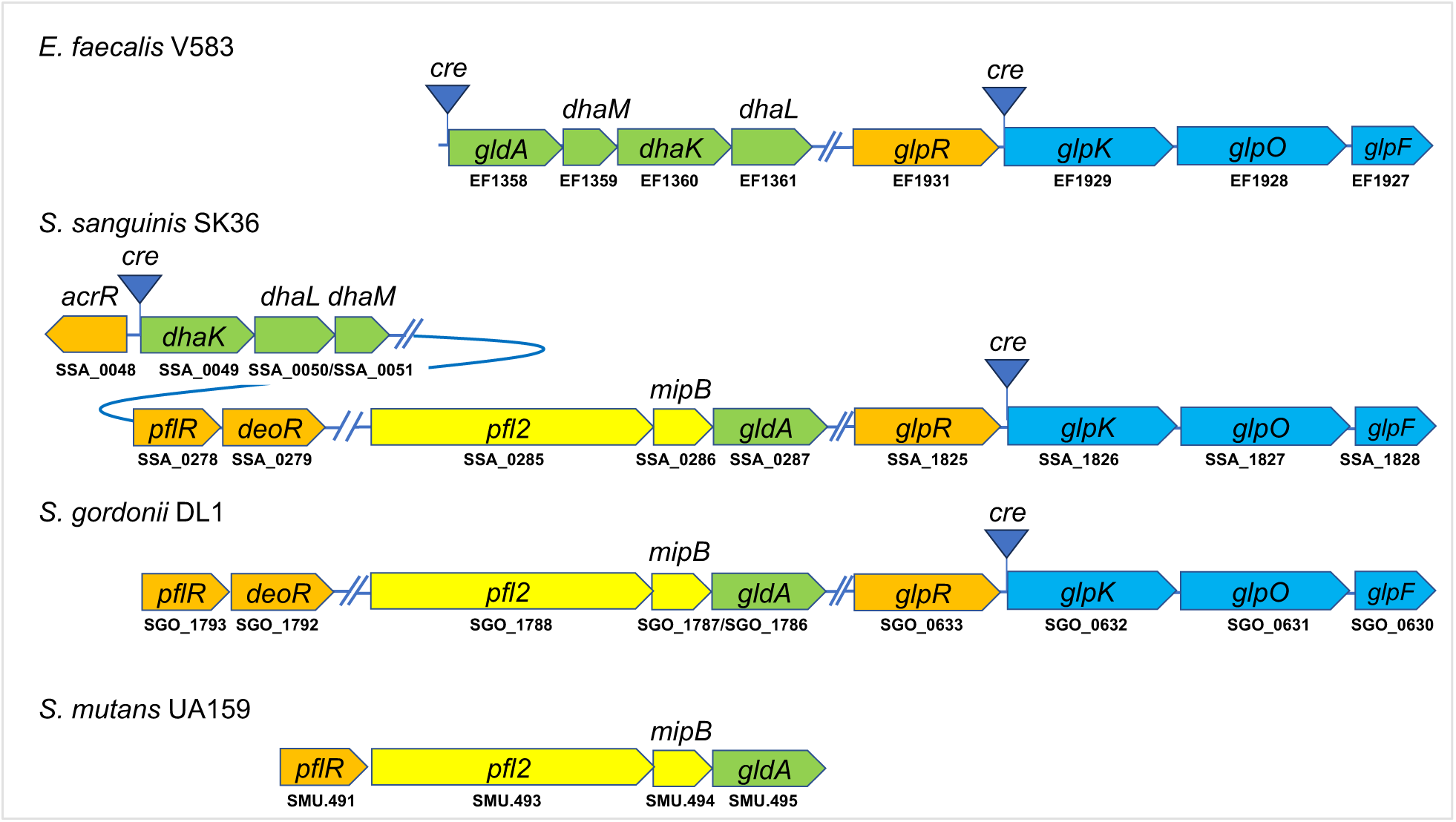
Diagram depicting genes required for glycerol metabolism in several lactic acid bacteria. Genes annotated for glycerol oxidation pathway *(glp)* are presented in blue whereas those for glycerol dehydrogenation pathway *(dha)* are in green. Putative transcription regulators associated with these genes are depicted in orange. Also denoted are *c*atabolite *r*esponse *e*lements *(cre)* located in the intergenic regions, a formate acetyltransferase *pfl2*, and a transaldolase-like protein *mipB*.

With glycerol being a secondary carbohydrate, transcription of the genes encoding both catabolic pathways is often under carbon catabolite repression (CCR)(19), with a CcpA (catabolite control protein A)-binding motif (*c*atabolite *r*esponse *e*lement, *cre*) being identified in regions upstream of these catabolic operons (*glp* and *dha*) in *B. subtilis* and various LAB species including *E. faecalis* and *S. sanguinis* (4)(Fig. 1). Another CCR mechanism exists to regulate the activity of the phosphorylation pathway at the enzymatic level, namely the requirement of GlpK for PTS-mediated phosphorylation by Enzyme I (EI) and phospho-carrier protein HPr at a conserved histidine residue for maximal activity (20, 21). These mechanisms suggest that metabolism of glycerol is likely repressed by the presence of preferred sugars such as glucose. Additional regulators have been reported to modulate glycerol metabolism, including the RNA-binding anti-terminator GlpP identified in *B. subtilis* that responds to Gly-3-P (9), and a positive regulator Ers in *E. faecalis* that regulates metabolism of glycerol, citrate, and arginine (22). A putative transcriptional regulator (*glpR,* Fig. 1) exists as part of the *glp* locus in enterococci (13) and most streptococci, whose function remains to be characterized.

The central role of carbohydrate metabolism in oral microbial ecology makes glycerol a novel subject when investigating metabolic interactions among constituents of the oral microbiome. For example, it is understood that oral commensals such as *Candida* (23) and *Corynebacterium* (17) can produce significant quantities (mM levels) of glycerol, and its metabolism and ecological implications remain largely unknown. We recently mapped the regulon of a glucose-PTS in *S. sanguinis* SK36 by deep RNA sequencing of a *manL* (EIIAB^Glc^) deletion mutant (24), a spontaneous mutant arisen due to enhanced fitness under acidic conditions in association with a reduced glucose transport (25). As part of the glucose-PTS regulon, genes of both glycerol metabolic pathways showed significantly higher expression in the *manL* mutant, especially the phosphorylation pathway (*glpKOF*) which showed the greatest increase (fold of change >70) within the entire transcriptome (24). Conversely, a recent RNA-seq analysis performed on a *ccpA* mutant of SK36 indicated that none of these glycerol catabolic genes was transcriptionally affected by loss of CcpA (26). The genome of *S. sanguinis* lacks the homolog of the anti-terminator GlpP found in *Bacillus*. Therefore, PTS could be regulating glycerol metabolism in *S. sanguinis* and related LAB in a CcpA-independent manner that significantly deviates from that in *Bacillus subtilis* (9, 10). Here we conducted a systematic study of glycerol catabolism by SK36, its regulation in response to other carbohydrates, and the implications for the ecology of the oral microbiome.

## Results

### Growth phenotypes of mutants deficient in glycerol metabolic enzymes and glucose-PTS

In contrast to previous findings suggesting a lack of glycerol fermentation by many lactic acid bacteria (12), recent research has shown significant effects of glycerol on the physiology of *S. sanguinis* in a manner dependent on the glycerol kinase (GlpK)(17). To understand the metabolism of glycerol and its effects on bacterial physiology, we constructed several genetic mutants (Fig. 1) in the background of *S. sanguinis* SK36 and tested their ability to grow in a synthetic medium (FMC) supplemented with glycerol as the sole carbohydrate (Fig. 2A). Despite the conservation of both *glpKOF* and the *gldA-dhaKLM* genes in the genome, SK36 could only produce low levels of growth that raised OD_600_ by about 0.05 units, a result consistent with previous findings (12). However, deletion of *glpK* or *glpR,* but not deletion of *glpF,* abolished this consistent, albeit low level of, growth. Complementation of *glpK* and *glpR* deletion each largely rescued the ability of the complemented strains to grow on glycerol (Fig. S1A). Compared to FMC supplemented with 20 mM glucose alone, the combination of glucose and glycerol (5 mM) did not significantly alter the growth for most strains. However, the *manL* mutant deficient in the major glucose-PTS permease (EIIAB^Man^) showed a significantly reduced growth rate and yield in FMC supplemented with glucose and glycerol combined in comparison to FMC containing only glucose; the *manL/glpK* double mutant did not (Fig. 2BC). These results suggested that genetic or biochemical constraints exist in *S. sanguinis* that limit the utilization of glycerol as a carbohydrate.

**Fig. 2.**
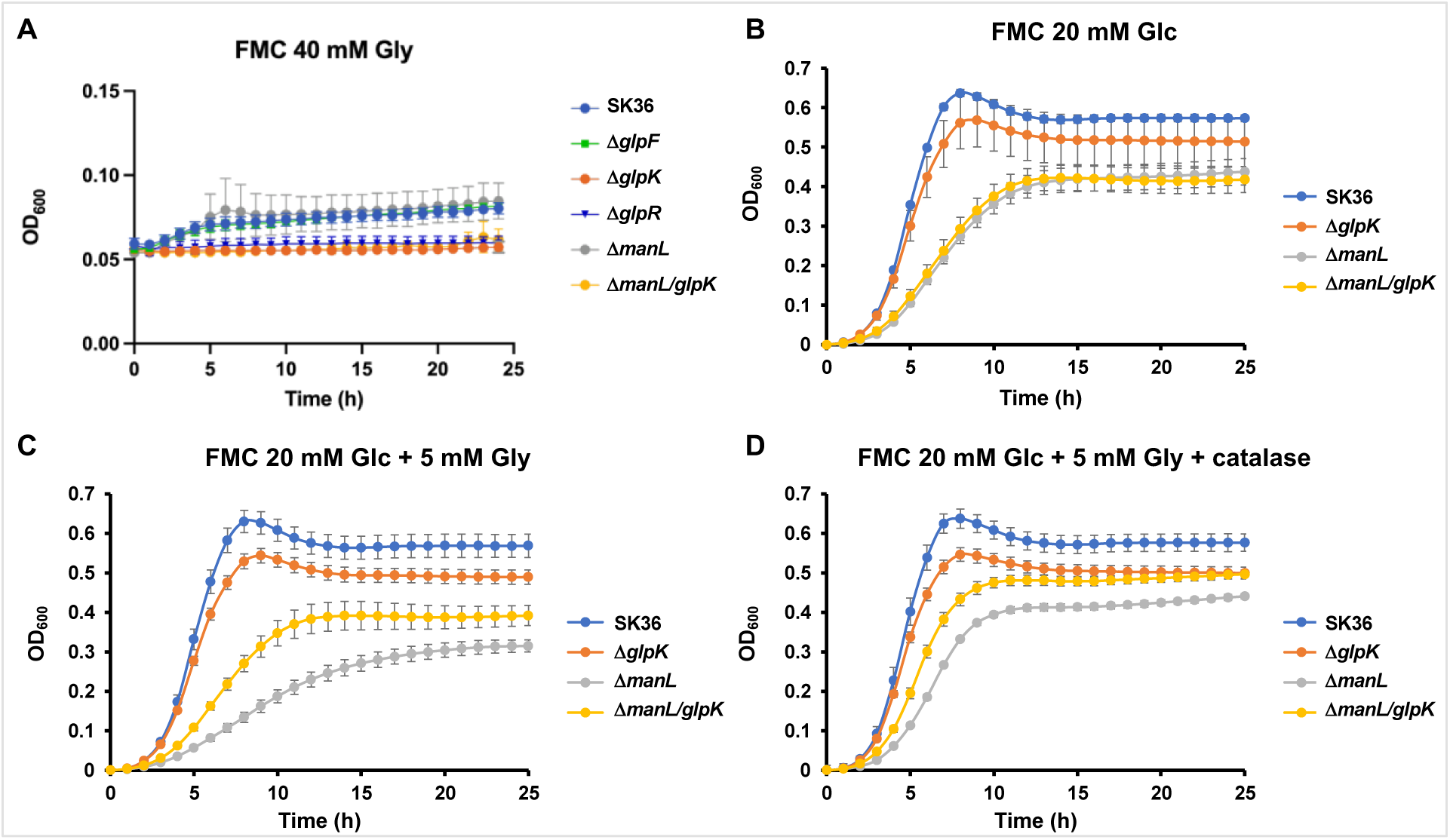
Growth curves measured in FMC medium modified to contain (A) glycerol, (B) glucose, (C) glucose and glycerol, and (D) glucose, glycerol and 5 μg/ml catalase. Strains were cultured to mid-exponential phase (OD_600_ = 0.5) in BHI, before being diluted 100-fold into 200 μl of defined medium (FMC) and loaded in a 96-well plate and covered with 60 μl mineral oil. OD_600_ was recorded using the Biotek Synergy 2 once every hour for 24 hours. The incubation was carried out aerobically at 37°C. Each sample was represented by at least 4 biological replicates and error bars denote standard deviations.

### Effects of glycerol metabolism on bacterial fitness through H_2_O_2_ production

To understand the cause of the reduced growth of the *manL* mutant in the presence of glycerol, we first measured the release of extracellular DNA (eDNA) in the culture medium as a simple way to assess bacterial autolysis (27). Elevated levels (by about 3-fold) of eDNA were noted in cultures of the *manL* mutant in the presence of 20 mM glucose and 5 mM glycerol compared to cultures prepared with only 20 mM glucose, yet no such difference was seen in the cultures of *manL/glpK* double mutant, the *glpK* mutant, or the wild-type parent (Fig. 3A). As glycerol-3-phosphate oxidation by the *glpKOF* pathway likely involves generation of H_2_O_2_, and H_2_O_2_ has been known to induce autolysis, we then measured H_2_O_2_ levels in the same culture media. The results showed levels of H_2_O_2_ in the supernatant of the SK36 cultures prepared with glucose and glycerol combined that were substantially higher compared to those prepared with glucose alone (increased from 1 to 1.2 mM, Fig. 3B), despite a lack of statistical significance. Importantly, the *manL* mutant which showed increased autolysis in the presence of glycerol, also produced significantly more H_2_O_2_ than the wild type in cultures supported by glucose (1.4 mM) and especially in cultures containing both glucose and glycerol (1.9 mM). In support of the notion that the *glp* pathway might have contributed to this increase in H_2_O_2_ levels, deletion of *glpK* in the *manL* background abolished the glycerol-dependent increase. Deletion of *glpK* alone in the SK36 background also resulted in the loss of response to the addition of glycerol. Finally, to confirm that differential production of H_2_O_2_ was the main cause of these growth and autolytic phenotypes, 5 μg/ml catalase was added to the culture medium containing a combination of glucose and glycerol. As shown in Fig 2D, relative to the same cultures prepared without catalase (Fig. 2C), addition of catalase largely rescued the growth of the *manL* mutant close to the level seen in FMC containing only glucose. Notably, addition of catalase also benefited growth of the *manL/glpK* double mutant.

**Fig. 3.**
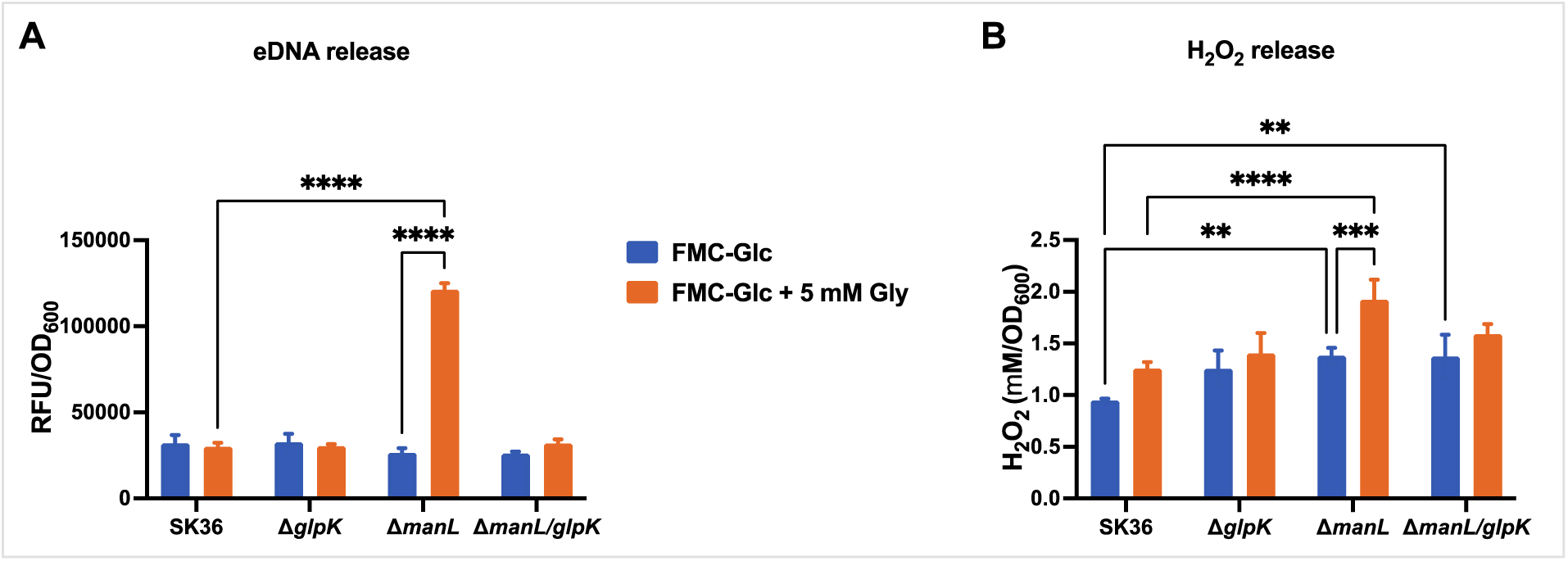
Glycerol induced in the *manL* mutant increased release of eDNA and H_2_O_2_. The bacteria were cultivated in FMC constituted with 20 mM glucose and 0 or 5 mM of glycerol for 24 h in an aerobic atmosphere with 5% CO_2_. The supernatant of each culture was harvested by centrifugation and subjected to measurement for (A) the relative levels of extracellular DNA by reacting with a fluorescent DNA dye, and (B) H_2_O_2_ concentrations using a biochemical reaction and a standard curve. At least three biological repeats were included in each sample, and the results were normalized against the cell density represented by OD_600_ of each bacterial culture. The bars represent the mean of the measurements and error bars denote the standard deviations. Asterisks indicate statistical significance assessed by two-way ANOVA followed by Tukey’s multiple comparisons test (*, *P* <0.05; **, *P* <0.01; ***, *P* <0.001; and ****, *P* <0.0001).

Therefore, H_2_O_2_ production from glycerol through the *glp* pathway is likely regulated by glucose and the glucose-PTS, the deletion of which resulted in production of H_2_O_2_ at levels that triggered excessive bacterial autolysis.

### Glycerol oxidation via GlpKO contributes to production of H_2_O_2_ independently of pyruvate oxidase

To further understand glycerol metabolism in *S. sanguinis* and the influence of the PTS, deletion mutants of *glpK*, *glpO*, *glpF, gldA,* and *glpR* were tested for their ability to catabolize glycerol along with mutants deficient in *manL, ccpA,* and *spxB*. We first carried out a Prussian Blue (PB) plate assay to assess the release of H_2_O_2_ by some of these strains on TY agar plates supplemented solely with glucose or glycerol. As indicated in Fig. 4A, SK36 produced significantly more H_2_O_2_ when incubated on plates containing glycerol than those containing glucose. Interestingly, while the *spxB* mutant largely lost the ability to produce H_2_O_2_ on glucose agar as expected, it continued to do so at levels comparable to the WT on plates containing solely glycerol. On the other hand, mutants deficient in *glpK,* or *glpO* (Fig. S2), produced significantly lower levels of H_2_O_2_ than the WT on glycerol agar plates, although they behaved much like the WT on glucose plates. Strain *glpKCom,* which had its *glpK* deletion corrected in the genome, produced WT levels of H_2_O_2_ on glycerol plates. These results suggested the ability of SK36 to produce H_2_O_2_ in a SpxB-independent, *glpKO*-dependent manner while catabolizing glycerol. Likely encoding a glycerol uptake facilitator protein, *glpF* is not essential to glycerol metabolism as its deletion did not result in a significant reduction in excretion of H_2_O_2_ on glycerol plates. In additional experiments (Fig. 4B) performed on glucose agar plates, mutants deficient in *ccpA* or *manL* (including *manL*/*glpK*) produced elevated levels of H_2_O_2_. This agrees with the theory that CcpA is the major regulator controlling transcription of *spxB* when cells are catabolizing glucose, where loss of ManL can alleviate CCR by reducing carbon flux (10). However, on glycerol agar plates, only the *manL* mutant produced higher levels of H_2_O_2_, and the *manL/glpK* double mutant behaved similarly to the *glpK* mutant, thus supporting the notion that glucose-PTS negatively regulates the *glp* pathway independently of CcpA.

**Fig. 4.**
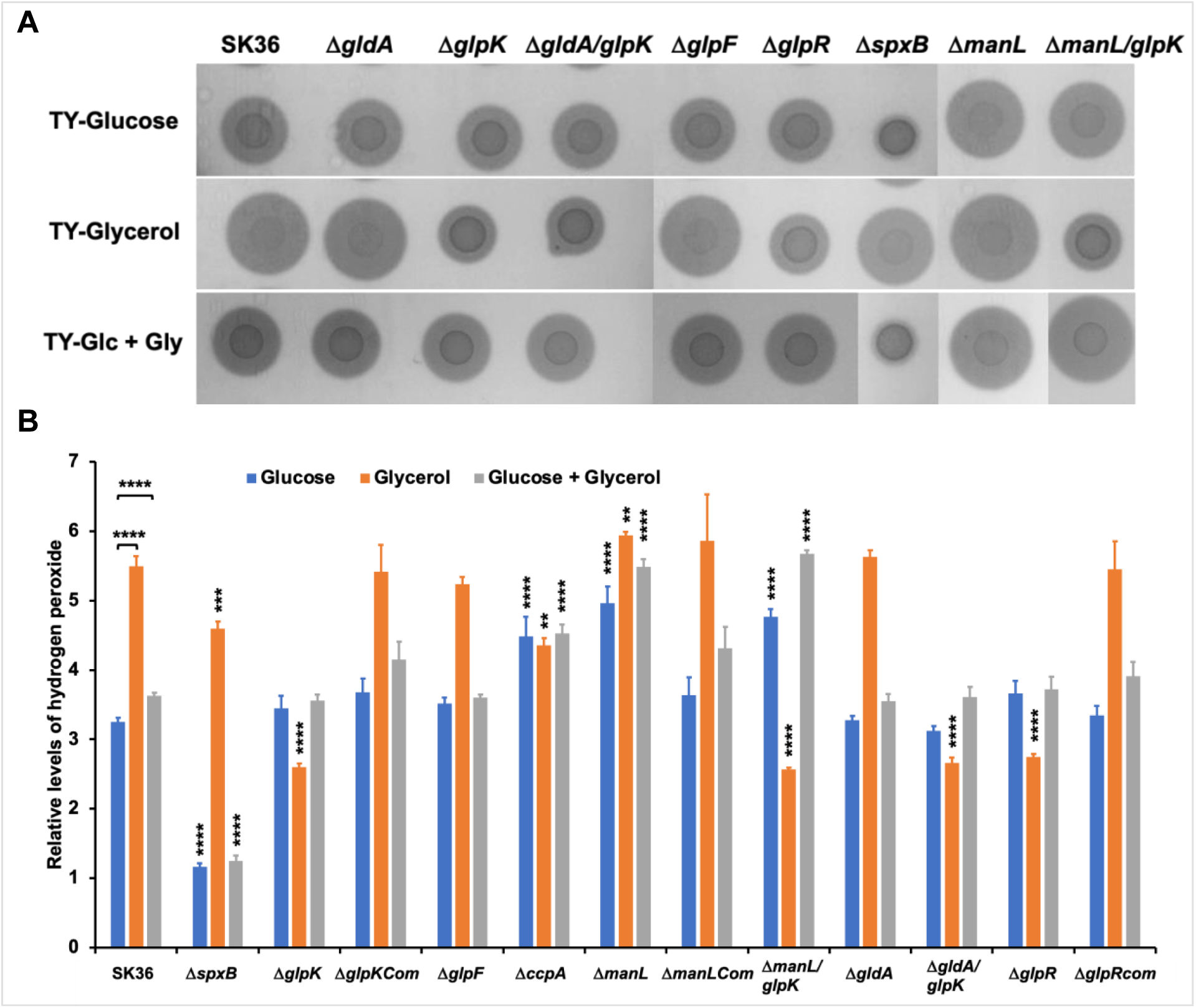
Prussian Blue (PB) plate assay for measurement of H_2_O_2_ release. Overnight cultures of bacteria in BHI were washed and dropped onto PB plates containing 20 mM glucose, 40 mM glycerol, or 20 mM glucose plus 5 mM glycerol. After another day of incubation in an ambient incubator maintained with 5% CO_2_, (A) the plates were photographed and (B) the width of the blue zone was measured to represent the relative amounts of H_2_O_2_ being released. At least three biological repeats were used for each sample and their average size of PB zone was used to plot the bar graph, with error bars representing standard deviations and asterisks denoting statistical significance relative to results of the parent strain SK36 assayed on the same carbohydrate (unless specified otherwise) (Student’s *t*-test; **, *P* <0.01; ***, *P* <0.001; and ****, *P* <0.0001).

In addition, strain Δ*gldA* did not show any discernable difference relative to the wild type in the production of H_2_O_2_ on either plate, and the *gldA/glpK* double mutant showed a phenotype similar to that of Δ*glpK.* Strain Δ*glpR*, on the other hand, behaved similarly to Δ*glpK* by not producing significant amounts of H_2_O_2_ on glycerol. Also, like Δ*glpKCom,* complementation of *glpR* rescued the production of H_2_O_2._ Considered together with the growth phenotype of Δ*glpR* on glycerol alone (Fig. 2), it is plausible that the *glpR* gene product is required for the expression of the *glp* operon.

We also tested the same strains using TY agar plates containing a combination of 20 mM glucose and 5 mM glycerol. In the presence of both carbohydrates, most of the strains tested generated a PB zone similar in size to that produced on glucose-only plates. However, several strains, including SK36, Δ*manL,* Δ*manL/glpK,* Δ*gldA/glpK* and Δ*glpRcom,* did consistently produce slightly more H_2_O_2_ than in TY agar containing glucose alone.

### Transcription analysis delineates the contributions of CcpA, PTS, and GlpR in regulating glycerol metabolism

A previous RNA-seq analysis of the *manL* mutant of *S. sanguinis* SK36 identified the *glp* operon genes as the most highly increased among the entire transcriptome (24) (Table S1). Also increased in expression in the *manL* mutant were the *dhaKLM* genes, though to a lesser degree. Unlike in *E. faecalis*, the first gene of the glycerol dehydrogenase pathway, *gldA* (Fig. 1) is encoded separately from the *dha* operon in *S. sanguinis*. To begin unravelling the influence of both glucose-PTS and CcpA on transcriptional regulation of glycerol metabolic genes, we performed qRT-PCR assays on several representative genes from both glycerol branches by first analyzing strains SK36, Δ*manL*, Δ*ccpA*, and Δ*manL/*Δ*ccpA* that were grown to the exponential phase with TY media supplemented with glucose, or a combination of glucose and glycerol.

We first compared all the cultures prepared with TY-glucose (Table 1). Deletion of *ccpA* resulted in a markedly higher expression by genes *dhaL* and *spxB*, whereas deletion of *manL* led to a similarly drastic increase in *glpK* mRNA levels. Deletion of *manL* also resulted in a significant increase, though to a lesser degree than in Δ*ccpA*, in expression of *dhaL* and *spxB*. By contrast, deletion of *ccpA* produced little to no change in *glpK* expression. Deletion of both *ccpA* and *manL* however resulted in the highest increase of expression by each of these three genes, although this further increase, relative to their changes in either single mutant, was modest (less than 2-fold). These results were consistent with the notion that CCR of *glpK,* and likely also *glpO* and *glpF,* is mediated by a mechanism independent of CcpA, whereas CCR of *dhaL* and *spxB* is mediated directly by CcpA, since the deficiency in glucose-PTS should also alleviate CCR mediated by CcpA (10). On the other hand, transcription of *gldA* appeared to be uncoupled from the rest of the dehydrogenation pathway, as it showed little change in the Δ*ccpA* background and moderately reduced transcription in the *manL* mutant, confirming the findings of the RNA-seq analyses in both Δ*ccpA* and Δ*manL* (24, 26). Last, deleting *glpR* resulted in little to no change to the transcript levels of all four genes analyzed in cultures grown with glucose alone.

**Table 1.** Relative abundance of mRNA levels in SK36 and various deletion mutants. Cultures (n = 3) of SK36 and isogenic mutants were prepared with TY or FMC containing 20 mM glucose (Glc), fructose (Fru), or galactose (Gal), with or without 5 mM glycerol (Gly). Asterisks denote statistical significance in mRNA abundance relative to results in SK36 grown under the glucose condition in the same type of medium (Assessed by Student’s *t*-test. *, *P* <0.05; **, *P* <0.01; ***, *P* <0.001; and ****, *P* <0.0001).

| Media |  | Strains | *gldA* | *glpK* | *dhaL* | *spxB* |
| --- | --- | --- | --- | --- | --- | --- |
| TY | Glc | SK36 | 1.01 | 1.01 | 1.00 | 1.01 |
| | | $\Delta glpR$ | 0.86 | 1.65 | 1.04 | 0.95 |
| | | $\Delta ccpA$ | 1.48 | 1.55 | 68.73**** | 10.25** |
| | | $\Delta manL$ | 0.45 | 65.50**** | 9.04** | 3.68* |
| | | $\Delta ccpA/\Delta manL$ | 1.12 | 79.09**** | 91.59**** | 13.43*** |
|  | Glc + Gly | SK36 | 1.46 | 1.93 | 0.90 | 1.53 |
| | | $\Delta glpR$ | 0.91 | 3.17* | 0.91 | 1.66 |
|  | Fru + Gly | SK36 | 2.30* | 0.76 | 1.11 | 0.36* |
| | | $\Delta manL$ | 2.11 | 0.64 | 1.03 | 0.38* |
|  | Gal | SK36 | 1.15 | 164.62**** | 31.68*** | 1.33 |
| | | $\Delta glpR$ | 1.76 | 2.09 | 46.06*** | 0.76 |
| | | $\Delta glpRCom$ | 0.65 | 218.96**** | 21.06*** | 3.60* |
|  | Gal + Gly | SK36 | 0.43 | 161.89**** | 47.82*** | 0.98 |
| | | $\Delta glpR$ | 0.88 | 2.37* | 44.37*** | 0.71 |
| | | $\Delta glpRCom$ | 0.65 | 218.59**** | 20.65*** | 3.08* |
| FMC | Glc | SK36 | 1.01 | 1.01 | 1.01 | 1.00 |
| | | $\Delta glpR$ | 1.36 | 1.58 | 1.23 | 0.88 |
|  | Glc + Gly | SK36 | 2.28* | 3.58* | 1.07 | 3.17* |
| | | $\Delta glpR$ | 2.02 | 5.84** | 0.82 | 4.53** |
|  | Gal | SK36 | 0.44* | 2.93* | 12.02*** | 17.42** |
| | | $\Delta glpR$ | 0.33* | 1.91 | 18.15*** | 8.77* |
|  | Gal + Gly | SK36 | 0.32** | 69.75*** | 5.44** | 12.58*** |
| | | $\Delta glpR$ | 0.29** | 2.55 | 14.55*** | 13.83*** |

To explore the influence of PTS other than glucose-PTS on glycerol metabolism, RT-qPCR was carried out using SK36 and Δ*manL* cultures prepared with TY containing a combination of 20 mM fructose and 5 mM glycerol. As the glucose-PTS does not transport fructose (25, 28), this experiment should determine if PTS EIIs other than EII^Man^ may trigger CCR on the *glp* operon. The results (Table 1) showed that just like glucose, the presence of fructose inhibited the expression of *glpK*—an effect not removed by the deletion of *manL*. Notably, compared to glucose, fructose showed no effect on *dhaL* expression yet reduced mRNA levels of *spxB* by close to 3-fold.

To understand the functions of GlpR, further transcriptional analysis was carried out in SK36 and Δ*glpR* cultivated in TY media supplemented with a combination of glucose and glycerol, galactose alone, or a combination of galactose and glycerol (Table 1). Addition of 5 mM glycerol to TY-glucose medium failed to induce in SK36 either the *glp* or *dha* pathway to a notable degree, a finding that is consistent with the strong CCR effect exerted by glucose. At the same time, deletion of *glpR* slightly increased expression of *glpK,* but not *dhaL,* in the presence of both glucose and glycerol. Several studies in streptococci have suggested that relative to glucose, catabolism of galactose via the tagatose pathway results in a minimum level of CCR since little CCR-inducing metabolites such as fructose-1,6-bisphosphate are produced (29–31). In SK36 cultures prepared with galactose alone, *glpK* showed the highest expression so far relative to cultures prepared with glucose, and *dhaL* also presented a considerable increase in mRNA levels. Deletion of *glpR* under galactose conditions abolished most of the increase seen with *glpK*, but not *dhaL*. This loss of *glpK* expression in Δ*glpR* was rescued by complementation in strain Δ*glpRCom*. These findings in galactose medium supported the notion of GlpR acting as an activator of the *glp* operon; however, they also suggested that glycerol may not be needed to induce the expression. When SK36 and Δ*glpR* were each cultured with a combination of galactose and glycerol, expression of both *glpK* and *dhaL* mirrored that in cultures prepared with galactose alone.

To rule out the possibility that TY media possesses at least low levels of glycerol, we repeated this experiment using the chemically defined medium FMC supplemented with galactose or a combination of galactose and glycerol. The results (Table 1) now showed that the expression of *glpK* in FMC constituted with galactose alone was only slightly higher than levels measured in FMC-glucose, however addition of glycerol to FMC-galactose drastically enhanced its expression. This result suggested that there likely was glycerol, or other inducing substrate, present in the TY-galactose medium at levels sufficient to induce the expression of *glpK*. Meanwhile, expression of *dhaL* increased in FMC-galactose even without addition of glycerol. We also repeated RT-qPCR using FMC supplemented with glucose +/-glycerol (Table 1), which largely supported findings made in TY media. Conversely, transcription of *gldA* in galactose-containing media was 2-to 3-fold lower than in glucose-based media. We subsequently measured glycerol levels in all the media available to us, including BHI, TY, TV, T-broth, FMC and BM (32), and found similar levels of glycerol (2-3 mM) in TY, TV and T-broth, and slightly lower amounts in BHI as well (Fig. S3). Considering this information, we have tried to conduct all the critical experiments using the FMC and BM and by washing cultures prepared with BHI before diluting into FMC or BM. Notably, we were not able to use FMC as the base of the Prussian Blue agar due to chemical incompatibility.

### Glycerol catabolism impacts growth on galactose by SK36 and by related oral streptococci

In light of the reduced CCR by galactose on glycerol catabolic genes, we conducted a growth analysis on strains SK36, Δ*glpR,* Δ*glpRCom*, Δ*glpK,* and Δ*glpKCom* by incubating them in FMC constituted with 20 mM galactose, with or without 5 mM of glycerol, in an aerobic environment supplemented with 5% CO_2_. While all strains grew relatively normally on galactose alone, addition of 5 mM glycerol markedly reduced the growth rate and yield of SK36, Δ*glpRCom*, and Δ*glpKCom* (Fig. 5AB). Interestingly, strains Δ*glpR* and Δ*glpK* each produced a growth in FMC containing a combination of galactose and glycerol that was comparable to growth in FMC with galactose alone (Fig. 5B). Considering the lack of CCR by galactose on expression of *glpK* in SK36 (Table 1), this GlpR- and GlpK-dependent growth reduction by glycerol can be explained as due to over production of H_2_O_2_, similar to the phenotype of the *manL* mutant in Fig. 2C. Indeed, when we added 5 μg/ml catalase to FMC containing a combination of galactose and glycerol, growth of SK36, Δ*glpRCom*, and Δ*glpKCom* was largely rescued to levels comparable to Δ*glpR* and Δ*glpK* (Fig. 5C). This finding not only demonstrated the drastic effect of *glp*-mediated H_2_O_2_ production on the physiology of SK36, but it also provided us a simple test to explore if similar response to glycerol existed in other oral streptococcus isolates.

**Fig. 5.**
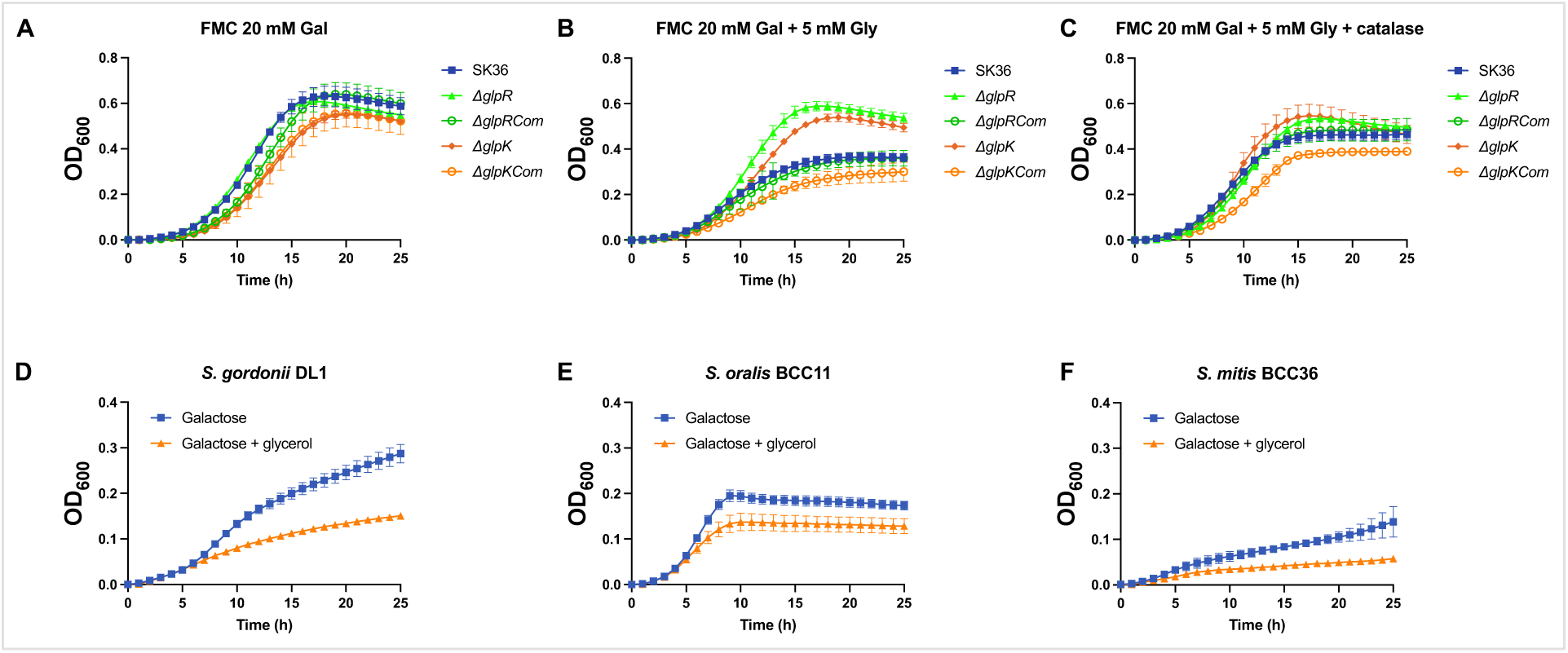
Growth curves constructed using *S. sanguinis* strains SK36 and its mutant derivatives (A, B, C), *S. gordonii* strain DL1 (D), *S. oralis* strain BCC11 (E), and *S. mitis* strain BCC36 (F). The media were based on FMC that was modified to contain 20 mM galactose, with or without 5 mM glycerol, and 5 μg/ml catalase (C). Each strain was represented by at least three biological repeats, with their means and standard deviations (error bars) in OD_600_ values being presented.

A BLAST search using the protein sequences of GlpK and GlpO identified respective homologs in most of the species belong to the mitis group streptococci. Two isolates each of *S. sanguinis, S. gordonii, S. oralis, S. cristatus,* and *S. mitis* were inoculated into FMC with galactose +/- glycerol as above. One strain each of *S. dentisani* and *S. intermedius* was also included as no homolog of GlpKO were identified in either species. Most of these strains have previously been analyzed by whole genome sequencing (33, 34). As shown in Fig. 5DEF (and Fig. S1D∼K), at least one strain each of *S. sanguinis, S. gordonii, S. oralis,* and *S. mitis* showed a reduced growth in the presence of glycerol relative to galactose-only condition. Some strains failed to grow on either medium, whereas *S. intermedius* BCC01 and two *S. cristatus* stains grew normally yet did not show a notable response to glycerol. Therefore, glycerol-mediated H_2_O_2_ production appears to be relatively well conserved in a group of streptococci that are associated with oral health, although further research is needed to confirm this conclusion.

### Influence of glycerol metabolism on streptococcal competition

We then performed plate and liquid-based competition assays to explore the effect of glycerol on antagonism between *S. sanguinis* and *S. mutans*. First, strains SK36, Δ*manL,* Δ*glpK,* and Δ*manL/glpK* were each inoculated on FMC-based agar plates supplemented with 20 mM glucose as the sole carbohydrate. FMC was selected in favor of TY to avoid contaminating glycerol and plates were incubated in an aerobic incubator with 5% CO_2_. *S. mutans* UA159 was inoculated to the right of *S. sanguinis* colonies after 24 h of incubation, followed by another day of incubation. Compared to the wild type SK36 (Fig. 6), deletion of *manL* resulted in a minor increase in inhibition of *S. mutans*. When 5 mM of glycerol was included in addition to glucose however, inhibition of *S. mutans* by strain Δ*manL* was significantly enhanced. At the same time, deletion of *glpK* in the background of *manL* reversed this effect, although deletion of *glpK* in the background of SK36 showed little impact. Interestingly, when glycerol was supplied as the sole carbohydrate, *S. sanguinis* but not *S. mutans* formed a visible colony on the agar plates (Fig. 6), echoing the limited growth of SK36 observed on glycerol alone (Fig. 2). When this experiment was repeated in galactose-based medium, with or without glycerol, SK36 and all its mutants greatly inhibited the growth of UA159 in comparison to UA159 inoculated on the plate alone, consistent with previous observation that both *spxB* and *glpK* were highly expressed under these conditions (Table 1) (35).

**Fig. 6.**
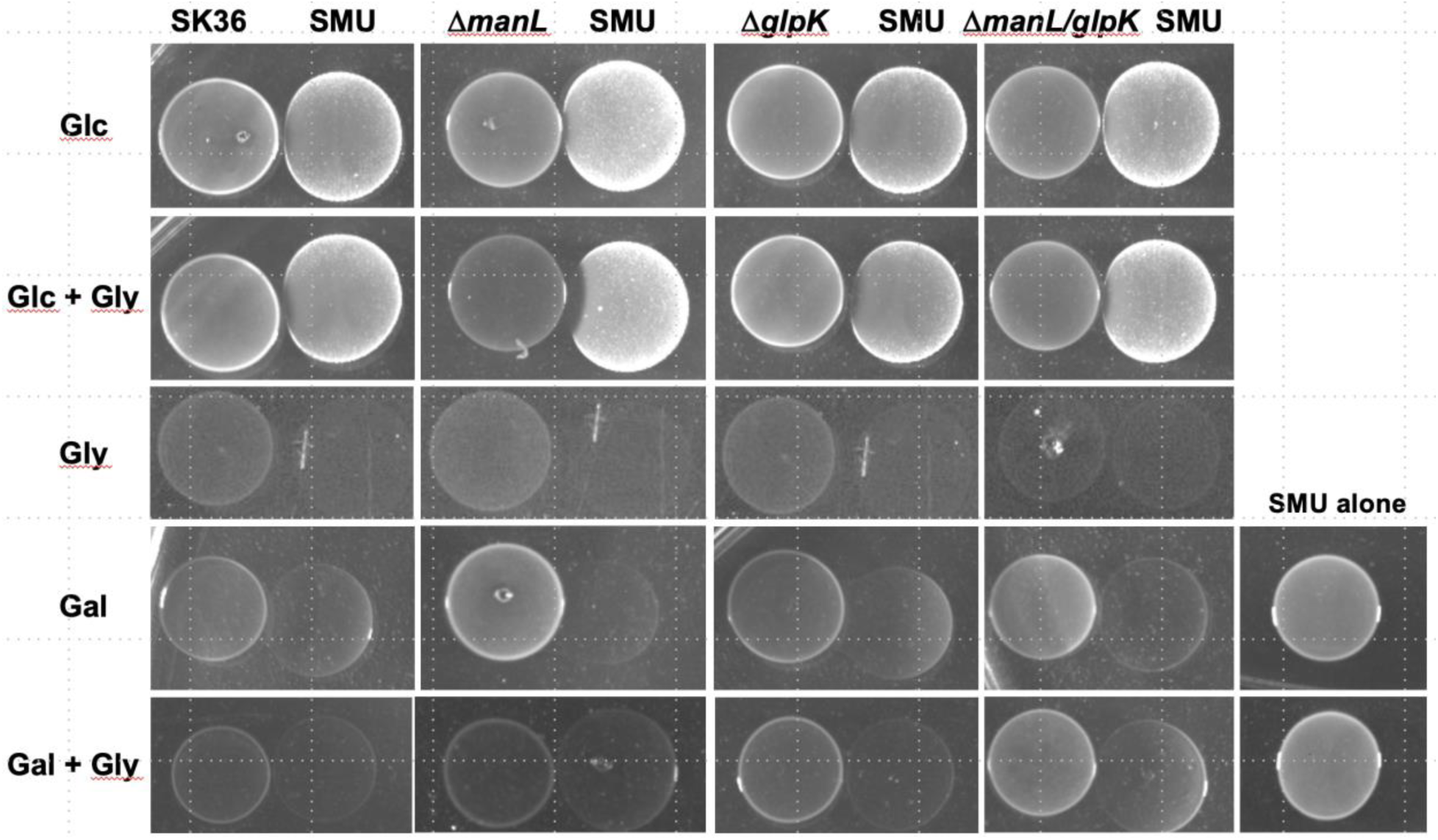
Deletion of *manL* enhanced antagonism of *S. mutans* in the presence of glycerol. To avoid contaminating carbohydrates, FMC was used as the base medium and was modified to contain 20 mM glucose, 20 mM glucose and 5 mM glycerol, 40 mM glycerol, 20 mM galactose, or 20 mM galactose and 5 mM glycerol. Each *S. sanguinis* strain was inoculated on the agar plates first. After an overnight of incubation, a fresh BHI culture of *S. mutans* UA159 (SMU) was inoculated to the immediate right of the colony, followed by another day of incubation before photography. Each interaction was tested at least twice on separate days, with a representative set of results being presented here.

Nonetheless, the *glpK* mutant did show less antagonism than SK36 in the presence of glycerol. These experiments confirmed glycerol metabolism by *S. sanguinis* (via function of GlpKO) as an ecologically significant source of H_2_O_2_, and the role of glucose-PTS in regulating this activity.

In consideration of the oxygen-limited condition in the oral cavity, a competition assay was carried out in planktonic cultures to mimic such a condition, by mixing *S. sanguinis* and *S. mutans* at approximately 1:1 ratio and cultivating overnight in an aerobic chamber maintained with 5% CO_2_. When these mixed cultures were diluted and allowed to compete in TY containing only 20 mM glucose, *S. mutans* UA159 was drastically more competitive over *S. sanguinis* SK36, with the competitive index (SSA/SMU) nearing 10^-8^. Little difference in the competition was noted when 5 mM glycerol was added to the medium that contained 20 mM glucose. However, when 40 mM glycerol was used as the only carbohydrate in TY medium, SK36 was approaching being equal in competitiveness with UA159, and this competitiveness was reduced by about 12-fold when the *glpK* gene was deleted (Fig. 7A). In each of these 20-h two-species cultures prepared with TY-glycerol, the total CFU was around 10^8^ to 10^9^ CFU/ml. Meanwhile, SK36, *glpK* and UA159 each yielded 10^8^ to 10^9^ CFU/ml viable cells after overnight incubation on TY-glycerol as single-species cultures. These results demonstrated the ability of glycerol to greatly enhance the competitive fitness of SK36 against UA159 mostly in the absence of glucose, a capacity that is not entirely dependent on an intact *glpK* gene. Interestingly, when the *gldA* mutant was tested similarly, it behaved much like the WT parent under all three conditions, showing no reduced competitiveness in TY containing glycerol.

**Fig. 7.**
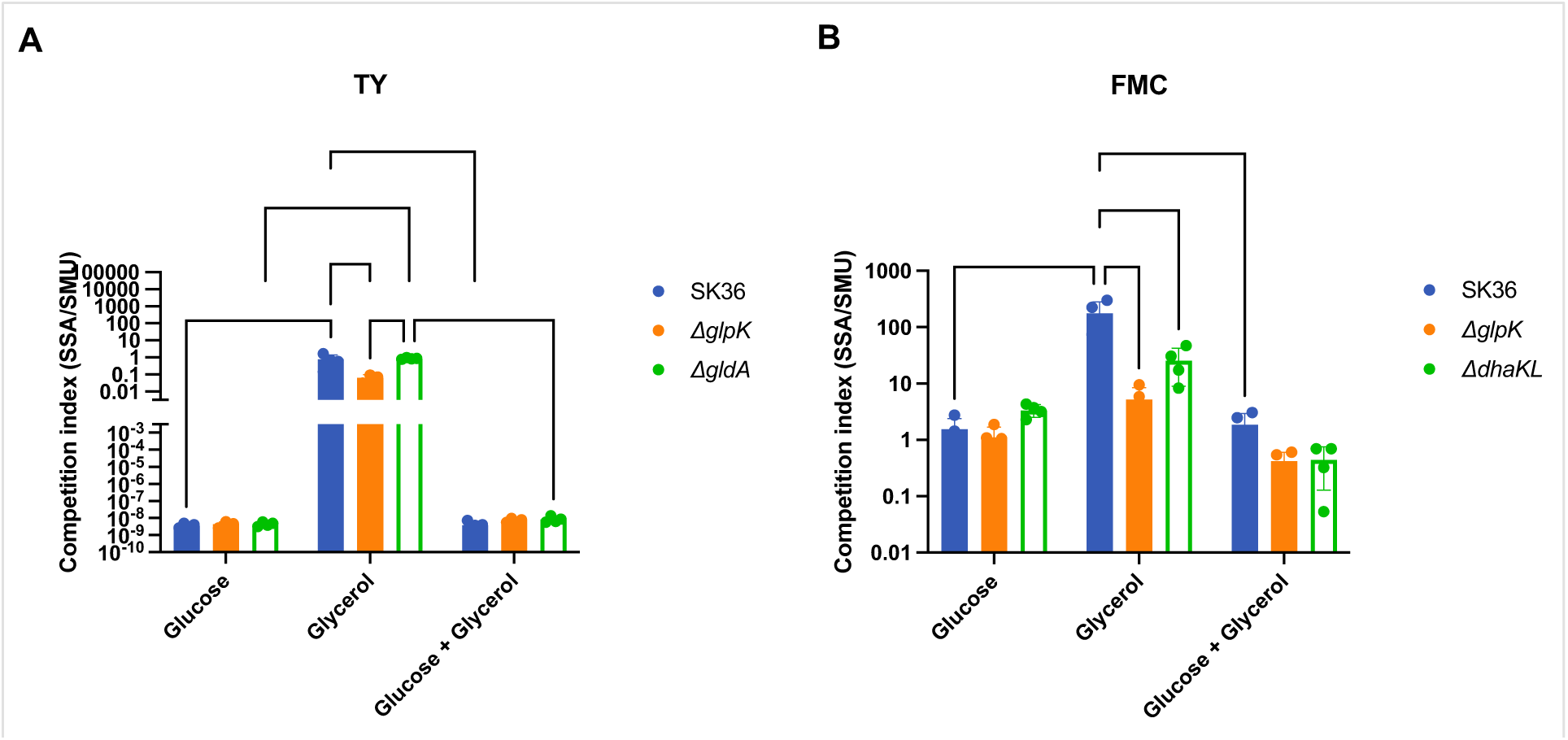
Glycerol greatly benefits *S. sanguinis* (SSA) against *S. mutans* (SMU) in a planktonic competition assay. Exponentially growing BHI cultures of *S. sanguinis* SK36, Δ*glpK,* Δ*gldA* and Δ*dhaKL* were each mixed with equal amounts of *S. mutans* strain UA159, inoculated at 100-fold dilution rate and allowed to grow overnight in TY (A) or FMC (B) medium supplemented with 20 mM glucose, 40 mM glycerol, or a combination of 20 mM glucose and 5 mM glycerol. CFU of each competing species before and after the competition was enumerated to calculate the competition indices, with values greater than 1 indicative of an advantage for *S. sanguinis*. Each strain was represented by 4 separate cultures, from which the means and standard deviations (error bars) were calculated and presented. Statistical analysis was carried out using Two-way ANOVA, followed by Tukey’s multiple comparisons (*, *P* <0.05; **, *P* <0.01; ***, *P* <0.001; and ****, *P* <0.0001).

Results obtained thus far suggested that the ORF annotated as *gldA* may not be contributing significantly to glycerol metabolism. Therefore, a new mutant deficient in both *dhaK* and *dhal,* Δ*dhaKL,* was constructed to represent the *dha* pathway. Strain Δ*dhaKL* showed no change in the ability to grow on glycerol (Fig. S1O) or produce H_2_O_2_ (Fig. S2) in comparison to its WT parent. We then repeated the planktonic competition assay by replacing Δ*gldA* with Δ*dhaKL,* and by using FMC as the base medium to avoid glycerol contamination. Overall, the *S. sanguinis* wild type was significantly more competitive when growing in FMC-based media than in TY-based media, likely owing to superior buffering capacity of the former. Relative to glucose, glycerol alone enhanced the competitiveness of *S. sanguinis* against *S. mutans* by 2 logs, and disruption of either the *glp* or *dha* pathway significantly reduced its competitiveness (Fig. 7B). Specifically, deletion of *glpK* resulted in a 34-fold reduction (*P* <0.0001) in competitive index in FMC-glycerol, as well as a 6-fold reduction in FMC containing glycerol and glucose combined, although the latter did not reach statistical significance. On the other hand, deletion of *dhaKL* also reduced the competitive index of SK36 by 7-fold on glycerol alone (*P* <0.0001) and by 3-fold on glycerol + glucose (*P* >0.05). Therefore, it appears that both the *glp* and *dha* branches, *glp* especially, are required for competitive fitness of *S. sanguinis* under planktonic condition in the presence of glycerol.

Last, the effect of glycerol on bacterial competition was tested in a biofilm setting. A dual-species biofilm model was established using *S. sanguinis* and *S. mutans* UA159 on the surface of a hydroxyapatite disc submerged in a BM medium (32) supplemented with 25% saliva, 2 mM sucrose, and 18 mM glucose. After 1 day of incubation, the medium of the biofilm was replaced with BM containing glucose, BM with glycerol, or BM base without any carbohydrate. 24 h later, the biofilm was harvested and CFUs of both species were enumerated. This experiment was meant to simulate a mature dental biofilm being exposed to a dose of glucose or glycerol. The results (Fig. 8) showed a strong negative effect of glycerol on the persistence of *S. mutans* by eliminating *S. mutans* cells from the biofilm. This effect of glycerol was dependent on the integrity of *glpK*, yet not that of *dhaKL*. When the same experiment was repeated in an anaerobic atmosphere, no significant effect was noted in association with glycerol (Fig. S4). Interestingly, deletion of *gldA* in SK36 resulted in a significant loss of CFU of *S. sanguinis* under all three conditions, including when treated with BM-glycerol where both bacteria returned no CFU from the biofilms. Furthermore, when different batches of pooled saliva were used in these assays, substantial variation was noted in the effectiveness of glycerol in inhibiting *S. mutans* in a *glpK*-dependent manner (Fig. S4). Considered together with the planktonic competition assay, both branches of the glycerol metabolism could benefit *S. sanguinis* during its interaction with *S. mutans,* with their relative contributions being dependent on experimental settings and/or certain environmental factors that are yet to be clarified.

**Fig. 8.**
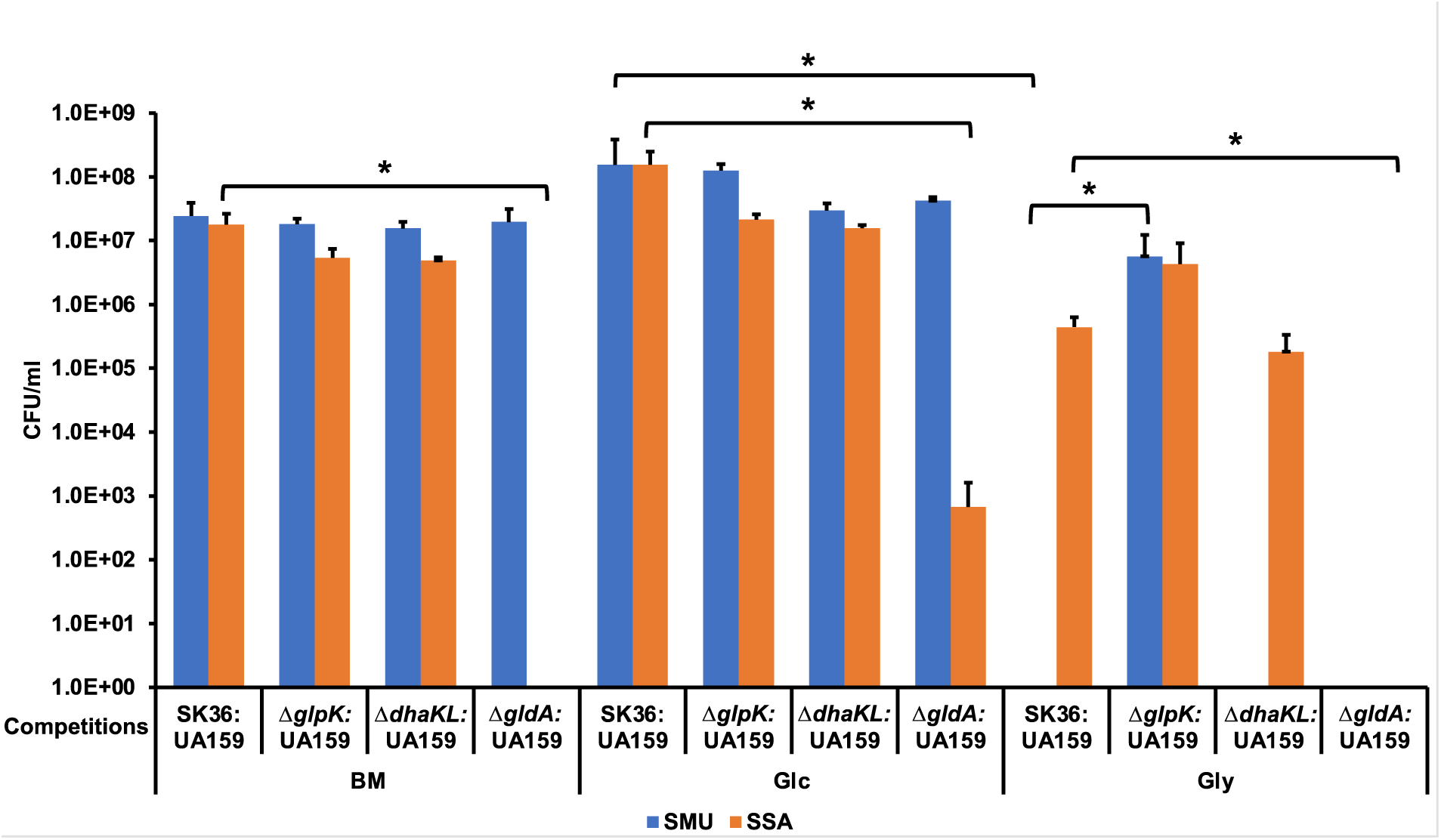
Glycerol promotes *S. sanguinis* (SSA) competition against *S. mutans* (SMU) in a two-species biofilm in a *glpK*-dependent manner. SK36 and its isogenic mutants Δ*glpK,* Δ*dhaKL,* and Δ*gldA* were each mixed with an equal volume of *S. mutans* UA159 culture and inoculated, at 1:100 ratio, into a BMGS medium held in a 24-well plate with a hydroxyapatite disc and kept in an aerobic atmosphere with 5% CO_2_. After 24 h of incubation, the culture supernatant was replaced with BM base, BM containing 18 mM glucose (Glc), or BM with 36 mM glycerol (Gly). After another day of incubation, the biofilm was washed and harvested by sonication, followed by serial dilution and CFU enumeration. Each strain was represented by 4 separate cultures and each condition 4 biofilm samples. The average CFU and standard deviation (error bars) of each species was used to plot the graph and for statistics (Students’ *t*-test; *, *P* <0.05).

### Expression of glycerol metabolic genes in dental plaque

To explore the contribution of glycerol to microbial homeostasis *in vivo*, a bioinformatic analysis was conducted using metatranscriptomic datasets generated previously in a deep sequencing project (24, 36). 70 human plaque samples were used in that study, including 34 caries-free (PF) and 36 caries-active samples, with the latter comprised of 11 samples from enamel lesions (PE) and 25 from dentin lesions (PD). After normalization for total reads per plaque, transcript abundance for individual genes was calculated and tabulated into pathways of known bacterial functions. Subsequent comparative analysis (Fig. 9) indicated an increase in overall transcript levels of most *S. sanguinis* pathways in PF samples relative to both caries-active groups. This was likely in large part due to the elevated abundance of *S. sanguinis* as a species in healthy biofilms compared to dysbiotic ones (37). We then identified two pathways associated with glycerol metabolism, one named “super-pathway for glycerol degradation to 1,3-propanediol” and the other “CDP-diacylglycerol biosynthesis pathway”, neither of which included the glycerol degradation pathways studied here (4). Nonetheless, as shown in Fig. 9, the “super-pathway for glycerol degradation to 1,3-propanediol” was among the top 10 functions in terms of fold change in PF vs PE samples, or PF vs PD samples. As a critical function in lipid metabolism (38), CDP-diacylglycerol biosynthetic genes also showed an above-average PF/PE ratio. We then analyzed transcript levels of specific genes of *glp* and *dha* pathways. Expression of the dehydrogenation pathway by *S. sanguinis* was above average levels in PF samples, with *gldA* at ∼2.6-fold that of the median of fold changes calculated over the entire genome, both against PE and PD samples (Table S2). Expression of *glpK* in PF samples was slightly above the median of the genome, at 1.3-fold, and so was a glycerophosphodiester phosphodiesterase (2-fold) with the purported function of hydrolyzing glycerophospholipid and liberating Gly-3-P. When we expanded the analysis to include other mitis group streptococci, a similar trend in expression of these glycerol metabolic genes was noted (Table S2), especially in the *Streptococcus_mitis_oralis_pneumoniae* subgroup, followed by *S. australis* and *S. gordonii*. Therefore, genes required for glycerol metabolism in *S. sanguinis* appeared to be actively expressed in dental plaque in association with oral health.

**Fig. 9.**
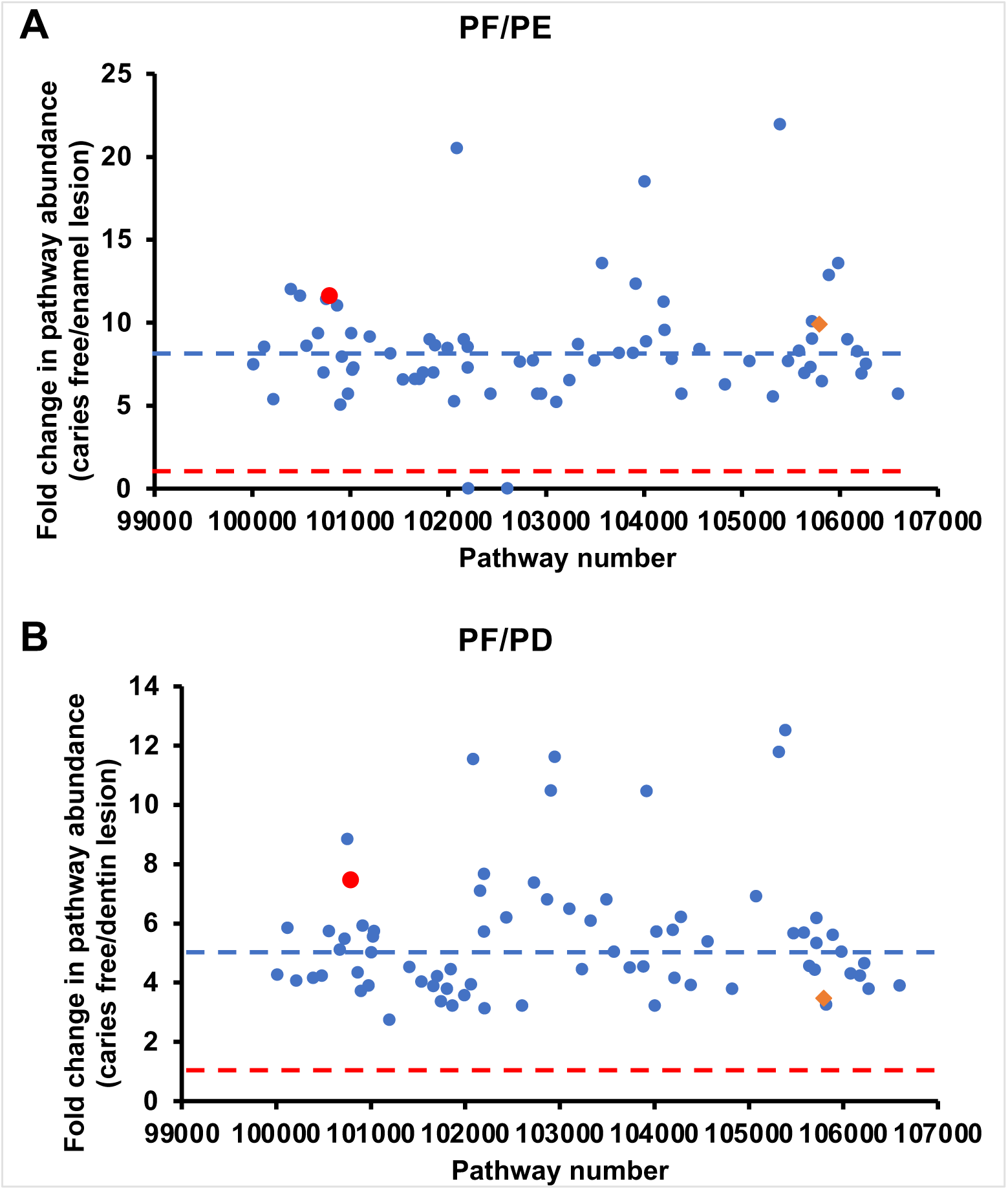
Glycerol metabolic pathway of *S. sanguinis* shows elevated expression in caries-free dental plaque samples. Metatranscriptomic analysis was performed on 70 plaque samples, 34 caries-free (PF) and 36 caries-active [11 from enamel sites (PE) and 25 from dentin sites (PD)]. Scatter plots show fold changes in transcript abundance in (A) caries-free vs enamel lesions and (B) caries-free vs dentin lesions. The red lines denote the value of 1 and the blue lines show the median of the population. The red circle represents the “super-pathway of glycerol degradation to 1,3-propanediol”, and the orange diamond represents the “CDP-diacylglycerol biosynthesis pathway”.

## Discussion

The main findings of this study included the following: i) While *S. sanguinis* SK36 cannot grow optimally on glycerol alone, a consistent, low-level of growth was achieved on mM levels of glycerol in a *glpK-, glpR*-dependent manner. ii) the *glp* pathway can produce H_2_O_2_ independently of SpxB by catabolizing glycerol in the presence of oxygen, and the level of H_2_O_2_ derived from glycerol readily resulted in both autolysis in *S. sanguinis* and inhibition of *S. mutans* in co-cultures. iii) Expression of the glycerol dehydrogenation (*dha*) branch is primarily regulated by CCR mediated by CcpA, whereas the glycerol phosphorylation branch (*glp*) requires GlpR and glycerol for induction and is subjected to CCR by the PTS. iv) Metabolism of glycerol enhanced the competitiveness of *S. sanguinis* against *S. mutans* primarily in a GlpK-dependent manner, especially when tested in a biofilm model. The *dha* branch also contributed to the persistence of *S. sanguinis* under either planktonic or biofilm setting. With glycerol being an abundant carbohydrate in the oral cavity and released by other microbiomes, these novel findings open the door to further exploring the influence of glycerol in human health and diseases.

Oral streptococci have long been considered incapable of catabolizing glycerol for growth or acid production (12). Our study on one hand partly confirmed these earlier findings, but on the other hand revealed a new dimension in the physiology of these abundant oral bacteria concerning their metabolic interactions with the rest of the microbiota. Since SK36 harbors all the known genes of both *glp* and *dha* branches, we can predict that their gene products can catabolize each glycerol molecule to obtain one molecule of pyruvate and one net gain of ATP, in addition to two NADH by way of dehydrogenation (*dha*) or one NADH and one H_2_O_2_ through the phosphorylation (*glp*) branch. One more ATP can be obtained if pyruvate is further oxidized, e.g., by SpxB, PFL or PDH branches, into acetate, although NADH/NAD^+^ ratio must be maintained either by NADH oxidase (Nox) or other pyruvate-reducing enzymes such as lactate dehydrogenases (LDH) or alcohol dehydrogenases (ADH). There are reasons to believe that the phosphorylation branch is favored by SK36 under most of our test conditions. According to RT-qPCR, the transcript levels of the *gldA* gene in SK36 grown with glucose were about half that of *glpK* and remained unchanged in most conditions or mutants we tested; expression of *dhaL* was not inducible by glycerol and largely independent of GlpR. Another supporting evidence is the substantial amounts of H_2_O_2_ produced by the *spxB* mutant on glycerol. If so, the amounts of pyruvate produced by the dehydrogenation branch are likely limited, and this inference is supported by the lack of substantial H_2_O_2_ yield by the *glpK* mutant on glycerol, with the *spxB* gene being intact (Fig. 4); and by the importance of *glpK* to interbacterial interaction as revealed in our dual-species competition assays (Fig. 7&8). Furthermore, *S. gordonii* which is closely related to *S. sanguinis* in glycerol phenotype (Fig. 5), lacks all three *dha* genes (Fig. 1). Nonetheless, results of the competition assays (Fig. 7&8) indicated that the dehydrogenation branch was indispensable in the competitive fitness of SK36, both with and without glycerol. Further research is needed to understand if environmental factors and additional genetic mechanisms contributed to such difference in our model systems.

Another question raised during our study was why glycerol alone cannot support SK36 to grow to the levels similar to glucose, especially since all necessary genes to support gluconeogenesis and pentose phosphate pathway are present in its genome (39). This remains unclear even as we try to exhaust the test conditions. Aside from the fact that gluconeogenesis is energetically costly and glycerol dehydrogenation appears constrained by *gldA-dhaKLM* expression patterns, production of H_2_O_2_ by glycerol phosphorylation could prove to be a double-edged sword. However, addition of catalase only resulted in a slight enhancement in growth by SK36 on glycerol as seen in Fig. 2A and Fig. S1M, whereas incubating it in an anaerobic chamber abolished all indications of any increase in optical density (Fig. S1LMN), arguing against the notion that either oxygen or production of H_2_O_2_ was the cause of the lack of substantial growth on glycerol. To appreciate the significance of this unique metabolic activity, perhaps it is more appropriate to consider it in the context of inter-microbial interactions. Recent work on interactions between *S. sanguinis* and *Corynebacterium durum*, an abundant oral commensal, has provided direct support to the availability of mM levels of glycerol from another bacterium and its effects on *S. sanguinis* (17). Another important source of glycerol is the commensal yeasts such as *Candida*. When we cultivated a *C. albicans* strain SC5314 in FMC supplemented with various carbohydrates, as much as 1 mM glycerol was detected in the spent media (Fig. S3). Research has shown that *C. albicans* significantly increases its activity of glycerol release when it transitions into a biofilm state (23), that *C. albicans* readily co-aggregates with commensal oral streptococci (40), and that mixing with *C. albicans* cultures for 30 minutes induced the transcription of glycerol-metabolic genes in *S. gordonii* (41). Given the dual role of *Candida* species as both commensals and pathobionts, we posit that bacterial metabolism of glycerol released by *Candida* could also impact health in a nuanced manner dependent on the genomic makeup of its microbial niche.

Like most secondary catabolic genes, both branches of glycerol metabolism are subjected to negative regulations by CCR. The novelty of our finding resides in the fact that CcpA appears to be directly controlling the transcription of only the *dha* branch whereas the PTS regulates the *glp* branch. Our initial findings suggest that glucose-PTS may be the chief mechanism controlling *glp* expression. However, as fructose similarly confers strong CCR on the same circuit (Table 1 and Fig. S1BC), it stands to reason that a mechanism shared by both glucose and fructose is likely responsible. Such phenotypes echo what have been reported in other streptococci, in particular *S. mutans*, where HPr has been suggested to work in concert with the glucose-PTS in controlling catabolic genes responsible for several secondary carbohydrate sources (42–44). The physiological significance of this bifurcated strategy in regulating two branches of the same metabolic pathway warrants detailed investigation, but as we have suggested previously concerning the regulation of a fructanase operon (*fruAB*) by both CcpA-dependent and PTS-dependent mechanisms, difference in the nature of the signals for these two systems allows for thresholded responses: PTS could recognize and respond to lower levels (tens of μM to low mM) of carbohydrates with specific structures, whereas CcpA primarily responds to changes in intracellular energy status triggered by higher amounts (>3 mM) of carbohydrates (45, 46). Glucose can be found in μM levels in saliva outside of mealtimes (47), whereas levels above mM can be expected in human blood or in the oral cavity during feeding. With GlpR being required for *glp* expression and the genome of SK36 apparently lacking an anti-terminator-style regulator (GlpP) that is required for CcpA-independent CCR of the *glp* pathway in *B. subtilis* (9), it seems reasonable to hypothesize that such a PTS-mediated regulation could be conducted via the ability of EI/HPr to phosphorylate and activate GlpK (17), whose product Gly-3-P in turn can stimulate GlpR’s DNA-binding activity. In addition, glycolytic intermediate fructose-1,6-bisphosphate has been reported to inhibit GlpK activities (48), which may also contribute to PTS-mediated regulation. Notably, we have preliminary evidence (Fig. 5) suggesting that additional peroxigenic streptococci likely also possess the ability to catabolize glycerol for H_2_O_2_ production, and similar operons containing *glpR* homologs appear to be conserved in at least some of these bacteria, including *S. pneumoniae* and *S. gordonii* (Fig. 1). It would be interesting to explore if similar metabolic strategies or CCR mechanisms exist in these related bacteria and if so, how that contributes to microbial homeostasis or host interactions in relevant microbiomes. Last, we cannot rule out the possibility that *glpK* is also subjected to the conventional CCR mediated by CcpA. After all, a near perfect *cre* sequence exists (Fig. S5) in the intergenic region upstream of the *glpK* sequence, and mutants defective in both *manL* and *ccpA* showed slightly higher *glpK* expression than that in either single mutant (Table 1). If so, CcpA could provide another layer of negative control on top of PTS on the catabolic operon, like the CCR of *fruAB* operon in *S. mutans* that involves both CcpA and the PTS (45).

It is perhaps worth noting that in most of our assays testing the functionality of the *glp* pathway, a *glpK* rather than a *glpO* genetic mutant was used. This was out of the concern that loss of glycerol oxidase GlpO alone, with the glycerol kinase GlpK being intact, would result in accumulation of Gly-3-P intracellularly which has been known to cause cytotoxicity effect (49). Given the presence of significant amounts of glycerol in several common media including BHI, the effects of Gly-3-P accumulation could incur a fitness cost and complicate interpretation of the data involving Δ*glpO*. Nonetheless, we have performed the PB plate assay on Δ*glpO* which showed a similar phenotype as that of Δ*glpK* (Fig. S2). Regarding the genetic uncoupling of *gldA* from the rest of the dehydrogenation branch, it could be the result of an evolution that favored H_2_O_2_ production by shunting glycerol toward the phosphorylation branch. A recent study in SK36 revealed two putative transcriptional regulators, SSA_0278 and SSA_0279, which were required for *gldA* expression in response to certain fatty acids during its interaction with *C. durum* (18), indicative of a possible specialization in function from or in addition to glycerol metabolism. The biofilm deficiency of strain Δ*gldA* revealed in Fig. 8 suggests certain novel functions of GldA in biofilm attachment by *S. sanguinis* that may or may not be related to catabolism of glycerol. Alternatively, there could exist a not-yet-identified isozyme that complements or replaces the function of GldA in dehydrogenating glycerol under certain conditions. Our finding that deletion of *dhaKL* genes, but not deletion of *gldA,* significantly impacted the fitness of SK36 in the presence of glycerol (Fig. 7) supports this notion. The gene product of *gldA* in *Escherichia coli* has been suggested to primarily catalyze the reverse reaction, by converting DHA into glycerol to avoid DHA-derived cytotoxicity; DHA or its metabolite being a reactive electrophile species capable of glycating various bio-molecules (50). In support of this notion, a likely transaldolase is encoded by an ORF (*mipB*) immediately upstream of *gldA* in SK36 (and in *S. gordonii* DL1), with the predicted function of converting DHA and glyceraldehyde 3-phosphate into fructose-6-phosphate. *S. mutans*, which lacks both *glp* and *dha* operons, also maintains the *gldA* and *mipB* homologs (Fig. 1) in its core genome (51) whose functions remain to be determined.

In conclusion, our research into glycerol metabolism by *S. sanguinis* revealed a novel pathway for generation of H_2_O_2_, an activity that appears uniquely significant regarding the competitiveness of the commensals and the health of the microbiome, rather than its contribution to bacterial growth. Further studies focusing on glycerol pathways in abundant oral bacteria, complex biofilms, and *in vivo* settings could help identifying potentially novel genetic mechanisms that can be targeted to enhance the homeostasis of oral microbiome and oral health.

## Materials and Methods

### Bacterial strains and culture conditions

Strains (Table 2) of *S. sanguinis* SK36 and its isogenic mutants, isolates of *S. mutans, S. gordonii,* and other oral streptococcal species were routinely maintained on BHI (Difco Laboratories, Detroit, MI) agar plates supplemented with 50 mM potassium phosphate (pH 7.2), and antibiotics (kanamycin at 1 mg/ml, erythromycin at 5 μg/ml) when necessary. Colonies were inoculated in liquid BHI medium and incubated overnight, before being dropped onto Tryptone-yeast extract medium (TY, 30 g of Tryptone and 5 g of yeast extract per liter) agar plates with or without Prussian Blue reagents (52), or agar plates based on synthetic FMC (53), or diluted into fresh TY or FMC medium that was modified to contain various specified carbohydrates. Unless specified otherwise, all cultures were incubated at 37°C in an aerobic atmosphere supplemented with 5% CO_2_.

**Table 2.** Strains used in this study.

| Strains | Relevant characteristics<sup>a</sup> | Source or reference |
| --- | --- | --- |
| SK36 | *S. sanguinis* wild type | Kitten laboratory |
| $\Delta glpK$ | SK36 *glpK::Em* | From SK36 |
| $\Delta glpKCom$ | $\Delta glpK$ complementation (*Km*) | From $\Delta glpK$ |
| $\Delta glpR$ | SK36 *glpR::Em* | From SK36 |
| $\Delta glpRCom$ | $\Delta glpR$ complementation (*Km*) | From $\Delta glpR$ |
| $\Delta glpF$ | SK36 *glpF::Em* | From SK36 |
| $\Delta gldA$ | SK36 *gldA::Em* | From SK36 |
| $\Delta gldA/glpK$ | SK36 *gldA::Em glpK::Km* | From $\Delta gldA$ |
| $\Delta dhaKL$ | SK36 *dhaKL::Em* | From SK36 |
| $\Delta spxB$ | SK36 *spxB::Em* | From SK36 |
| $\Delta manL/glpK$ | SK36 *manL::Km glpK::Em* | From $\Delta manL$ |
| MMZ1945 | SK36 *gtfP::Em* | (24) |
| $\Delta manL$ | SK36 *manL::Km* | (25) |
| MMZ1905 | SK36 *manLComp::Em* | (25) |
| MMZ1913 | SK36 *ccpA::Km* | (25) |
| MMZ1910 | SK36 *ccpA::Em manL::Km* | (25) |
| UA159 | *S. mutans* wild type, *perR*<sup>+</sup> | ATCC 700610 |
| MMZ1939 | UA159/pDL278::*gfp* (*Sp*) | From UA159 |

1023 <sup>a</sup> *Km*, kanamycin resistance cassette; *Em*, erythromycin; *Sp*, spectinomycin.

### Construction of genetic mutants

Genetic mutations were engineered by following a protocol based on allelic exchange using mutator DNA molecules generated using PCR amplification followed by ligation by Gibson Assembly (25). Briefly, two homologous DNA fragments, DNA_1 and DNA_2, flanking the target gene *X* were amplified using primers engineered to contain short DNA sequences that allowed an overlap between DNA_1 and the 5’ of the antibiotic marker, and an overlap between the 3’ of the antibiotic marker and DNA_2. These three DNA fragments were subsequently fused together in the same Gibson Assembly reaction to yield the mutator DNA. Wild-type *S. sanguinis* strain SK36 was cultured in BHI till early exponential phase (OD_600_ = 0.05-0.08), when competence stimulating peptide (synthesized at ICBR protein core, University of Florida) and the mutator DNA were added to the culture, followed by 3 h of incubation before plating on selective agar plates containing specific antibiotics (25). Colonies obtained this way were validated by PCR coupled with Sanger sequencing targeting the region of mutagenesis. All DNA oligos (listed in Table S3) were synthesized by Integrated DNA technologies (Coralville, IA).

Genetic complementation was carried out using a “knock-in” strategy detailed in our previous publication (25). Briefly, to correct the allelic replacement in the deletion mutant of gene X (e.g., Δ*X::Em*), the upstream flanking fragment DNA_1 was extended by using a different primer (Comp-2GA, Table S3) to include the entire coding sequence of the wild-type gene *X*. A mutator DNA was then created by ligating together this new DNA_1 fragment, the original downstream flanking fragment DNA_2, and an alternative antibiotic marker (e.g., *Km*). After transforming Δ*X::Em* with this new mutator DNA, Km-resistant colonies were picked, confirmed for sensitivity to erythromycin, and validated by sequencing similarly as above.

### Measurement of relative abundance of eDNA in bacterial cultures

Concentrations of extracellular DNA (eDNA) in FMC culture supernatants were assessed by mixing them with a fluorescent DNA dye, SYTOX Green (Ex/Em 504/523 nm; Thermo Fisher Scientific, Waltham, MA), and reading the intensity of light emission at 523 nm on a 96-well, Synergy H1 multimode plate reader (Agilent Technologies, Santa Clara, CA). The relative abundance of eDNA was presented as RFU normalized against OD_600_ of the bacterial culture. The validity of this assay was confirmed using a DNA standard curve as detailed in our previous publication (27).

### Measurement of H_2_O_2_ released by bacterial cultures

Prussian Blue (PB) plate assay: Each bacterial strain was cultivated overnight in liquid BHI medium, harvested by centrifugation, washed and resuspended with sterile PBS. 10 μl of the cell suspension was then inoculated onto TY agar plates supplemented with specified carbohydrates. After overnight incubation in an aerobic incubator containing 5% CO_2_, the plates were photographed, and the width of the PB zone was quantified using ImageJ software.

Measurement of H_2_O_2_ levels in liquid cultures was carried out using an enzymatic assay according to a previously published protocol detailed elsewhere (34).

### Plate-based antagonism assay

*S. sanguinis* strains were cultivated till exponential phase (OD_600_ = 0.5), from which a 5-μl aliquot was placed on an FMC agar plate modified to contain specified carbohydrates and incubated for 24 h to form a colony. An overnight culture of *S. mutans* UA159 prepared in BHI was then inoculated to the right of the colony. The plates were incubated for another day before being photographed.

### Mixed-species competition in planktonic cultures

For interspecies competition in liquid cultures, differentially marked strains of *S. sanguinis* (Em, MMZ1945) and *S. mutans* (Sp, MMZ1939) were each cultured in BHI to the exponential phase (OD_600_ = 0.5). An inoculum of approximately 10^6^ CFU/ml of SK36, or its otherwise isogenic mutants, together with an inoculum of similar numbers of *S. mutans* were added to a TY medium or FMC supplemented with 20 mM glucose, 40 mM glycerol, or a combination of 20 mM glucose and 5 mM glycerol, then cultured for 20 h in a 5%-CO_2_ environment at 37°C. At both the start and the end of the experiment, cultures were sonicated for 15 sec, serially diluted and plated on respective antibiotic plates to enumerate CFU of both species. The competitive index was calculated using the following formula: [SSA(*t*_end_)/SMU(*t*_end_)] / [SSA(*t*_start_)/SMU(*t*_start_)], with values >1 indicating SSA (*S. sanguinis*) being more competitive than SMU (*S. mutans*), and vice versa.

### Dual-species biofilm assay

Pooled donor (n >4) saliva was collected according to IRB protocol (University of Florida IRB201500497), heat inactivated by incubating at 60°C for 60 min, clarified by centrifugation, and filter sterilized. 25% (v/v) of saliva was then added to a biofilm base medium (BM) (32) supplemented with 2 mM sucrose and 18 mM glucose (BMGS). 500 μl of BMGS and a sterile hydroxyapatite (HA) disc was placed in each well of a 24-well microtiter plate. Differentially marked *S. sanguinis* (*Em*) and *S. mutans* (*Sp*) strains were each cultured to the exponential phase and diluted at 100-fold into the microtiter plate, followed by incubation at 37°C in an anaerobic chamber supplied with 5% CO_2,_ 10% H_2_ and 85% N_2_, or in an aerobic incubator maintained with 5% CO_2_. After 24 h, the culture supernatant was removed gently and replaced with BM-glucose (18 mM), BM-glycerol (36 mM), or BM with no carbohydrate, followed by another day of incubation in the same environment. Subsequently, the HA discs were removed and washed three times by dipping in a sterile PBS, from which bacterial biomass was dispersed into 1 ml PBS by sonication (FB120 water bath sonicator, Fisher Scientific) at 100% power, twice, for 15 sec. Each sample was then serially diluted and plated onto selective BHI agar plates for CFU enumeration. Each strain was represented by 4 biological repeats and each condition by 4 biofilm samples.

### RNA extraction and RT-qPCR

Bacterial strains were each inoculated into BHI for an overnight culture, which was diluted 20-fold into TY or FMC supplemented with carbohydrates as specified. To eliminate glycerol contamination, the overnight BHI cultures were collected by centrifugation and washed twice with sterile PBS before being diluted into the FMC media. Cells were harvested when the OD_600_ reached 0.5∼0.6, from which total RNA was extracted by following an established protocol using the RNeasy mini-kit and an RNase-free DNase I solution (Qiagen, Germantown, MD) for in-column gDNA removal (24). The total RNA was used in generation of cDNA using a reverse transcription kit (Bio-Rad, Hercules, CA) with gene-specific anti-sense primers, followed by quantitative PCR analysis using the CFX96 system and SYBR Green Supermix (Bio-Rad). The relative abundance of mRNA levels of each target gene was calculated using a ΔΔC_t_ method relative to a house-keeping gene (*gyrA*) used as internal control.

### Bioinformatic analysis

All bioinformatics was carried out using the HiperGator cluster computer at the University of Florida. On all paired-end reads across all samples, quality control of sequencing data was performed with FASTQC (54), and adapter contamination and quality trimming (cutoff = 30) were done with Cutadapt (55). Subsequently, reads were mapped against reference genomes via BWA-mem (56) by removing those derived from human host (GRCh38) and 16S rRNA (Silva database) (57). We used Samtools (58) and FASTQ for handling and sorting of BAM files, and Bedtools (59) for recovery of unmapped reads. Finally, pathways profiling for each dataset was carried out using the Humann2 (60) pipeline, and the subsequent differential expression analysis was done in R Statistical Language (61) with base and edgeR (62) packages.

### Statistical analysis and data availability

Statistical analysis of the data was carried out using the software of Prism (GraphPad of Dotmatics, San Diego, CA). Any data, strains, and materials generated by this study will be available upon request from the authors for research or validation purposes.

## Supporting information

Includes Figures S1 to S5 and Tables S1 and S3.

## Acknowledgments

This work was supported by a grant DE12236 from NIDCR and a startup fund to LZ from University of Florida. PC was an undergraduate visiting scholar supported in part by a scholarship from Nankai University, Tianjin, China. ZAT was supported in part by a T90 training grant DE021990 from NIDCR. We appreciate the support from Dr. Robert Burne in granting us access to the metatranscriptomic datasets of their probiotic study.

## Reference

1. Semkiv MV, Ruchala J, Dmytruk KV, Sibirny AA. 2020. 100 Years Later, What Is New in Glycerol Bioproduction? Trends Biotechnol 38:907–916.

2. Nevoigt E, Stahl U. 1997. Osmoregulation and glycerol metabolism in the yeast *Saccharomyces cerevisiae*. FEMS Microbiol Rev 21:231–241.

3. Blötz C, Stülke J. 2017. Glycerol metabolism and its implication in virulence in Mycoplasma. FEMS Microbiol Rev 41:640–652.

4. Doi Y. 2019. Glycerol metabolism and its regulation in lactic acid bacteria. Appl Microbiol Biotechnol 103:5079–5093.

5. Hames C, Halbedel S, Hoppert M, Frey J, Stülke J. 2009. Glycerol metabolism is important for cytotoxicity of *Mycoplasma pneumoniae*. J Bacteriol 191:747–53.

6. Joseph B, Mertins S, Stoll R, Schar J, Umesha KR, Luo Q, Muller-Altrock S, Goebel W. 2008. Glycerol metabolism and PrfA activity in *Listeria monocytogenes*. J Bacteriol 190:5412–30.

7. Staerck C, Wasselin V, Budin-Verneuil A, Rincé I, Cacaci M, Weigel M, Giraud C, Hain T, Hartke A, Riboulet-Bisson E. 2021. Analysis of glycerol and dihydroxyacetone metabolism in *Enterococcus faecium*. FEMS Microbiol Lett 368.

8. Muller C, Cacaci M, Sauvageot N, Sanguinetti M, Rattei T, Eder T, Giard J-C, Kalinowski J, Hain T, Hartke A. 2015. The intraperitoneal transcriptome of the opportunistic pathogen *Enterococcus faecalis* in mice. PLOS ONE 10:e0126143.

9. Darbon E, Servant P, Poncet S, Deutscher J. 2002. Antitermination by GlpP, catabolite repression via CcpA and inducer exclusion triggered by P-GlpK dephosphorylation control Bacillus subtilis glpFK expression. Mol Microbiol 43:1039–52.

10. Deutscher J, Francke C, Postma PW. 2006. How phosphotransferase system-related protein phosphorylation regulates carbohydrate metabolism in bacteria. Microbiol Mol Biol Rev 70:939–1031.

11. García-Solache M, Rice LB. 2019. The Enterococcus: a Model of Adaptability to Its Environment. Clinical Microbiology Reviews 32:10.1128/cmr.00058-18.

12. Gunsalus IC, Sherman JM. 1943. The fermentation of glycerol by streptococci. J Bacteriol 45:155–62.

13. Bizzini A, Zhao C, Budin-Verneuil A, Sauvageot N, Giard J-C, Auffray Y, Hartke A. 2010. Glycerol Is Metabolized in a Complex and Strain-Dependent Manner in *Enterococcus faecalis*. J Bacteriol 192:779–785.

14. Postma PW, Lengeler JW, Jacobson GR. 1993. Phosphoenolpyruvate:carbohydrate phosphotransferase systems of bacteria. Microbiol Rev 57:543–94.

15. Doi Y. 2018. Lactic acid fermentation is the main aerobic metabolic pathway in Enterococcus faecalis metabolizing a high concentration of glycerol. Applied Microbiology and Biotechnology 102:10183–10192.

16. Kilian M, Mikkelsen L, Henrichsen J. 1989. Taxonomic study of viridans streptococci: description of *Streptococcus gordonii* sp. nov. and emended descriptions of *Streptococcus sanguis* (White and Niven 1946), *Streptococcus oralis* (Bridge and Sneath 1982), and *Streptococcus mitis* (Andrewes and Horder 1906). IJSEM 39:471–484.

17. Treerat P, Anderson D, Giacaman RA, Merritt J, Kreth J. 2023. Glycerol metabolism supports oral commensal interactions. The ISME Journal 17:1116–1127.

18. Treerat P, Redanz U, Redanz S, Giacaman RA, Merritt J, Kreth J. 2020. Synergism between *Corynebacterium* and *Streptococcus sanguinis* reveals new interactions between oral commensals. Isme j 14:1154–1169.

19. Deutscher J. 2008. The mechanisms of carbon catabolite repression in bacteria. Curr Opin Microbiol 11:87–93.

20. Charrier V, Buckley E, Parsonage D, Galinier A, Darbon E, Jaquinod M, Forest E, Deutscher J, Claiborne A. 1997. Cloning and Sequencing of two Enterococcal glpK Genes and Regulation of the Encoded Glycerol Kinases by Phosphoenolpyruvate-dependent, Phosphotransferase System-catalyzed Phosphorylation of a Single Histidyl Residue*. Journal of Biological Chemistry 272:14166–14174.

21. Deutscher J, Bauer B, Sauerwald H. 1993. Regulation of glycerol metabolism in *Enterococcus faecalis* by phosphoenolpyruvate-dependent phosphorylation of glycerol kinase catalyzed by enzyme I and HPr of the phosphotransferase system. J Bacteriol 175:3730–3.

22. Riboulet-Bisson E, Hartke A, Auffray Y, Giard JC. 2009. Ers controls glycerol metabolism in *Enterococcus faecalis*. Curr Microbiol 58:201–4.

23. Desai JV, Bruno VM, Ganguly S, Stamper RJ, Mitchell KF, Solis N, Hill EM, Xu W, Filler SG, Andes DR, Fanning S, Lanni F, Mitchell AP. 2013. Regulatory role of glycerol in *Candida albicans* biofilm formation. mBio 4:e00637–12.

24. Zeng L, Walker AR, Burne RA, Taylor ZA. 2022. Glucose phosphotransferase system modulates pyruvate metabolism, bacterial fitness, and microbial ecology in oral streptococci. J Bacteriol 205:e0035222.

25. Zeng L, Walker AR, Lee K, Taylor ZA, Burne RA. 2021. Spontaneous mutants of *Streptococcus sanguinis* with defects in the glucose-PTS show enhanced post-exponential phase fitness. J Bacteriol 203:JB0037521.

26. Bai Y, Shang M, Xu M, Wu A, Sun L, Zheng L. 2019. Transcriptome, phenotypic, and virulence analysis of *Streptococcus sanguinis* SK36 wild type and its CcpA-null derivative (ΔCcpA). Front Cell Infect Microbiol 9:411.

27. Zeng L, Burne RA. 2019. Essential roles of the *sppRA* fructose-phosphate phosphohydrolase operon in carbohydrate metabolism and virulence expression by *Streptococcus mutans*. J Bacteriol 201:e00586–18.

28. Abranches J, Chen YY, Burne RA. 2003. Characterization of *Streptococcus mutans* strains deficient in EIIAB^Man^ of the sugar phosphotransferase system. Appl Environ Microbiol 69:4760–9.

29. Zeng L, Das S, Burne RA. 2010. Utilization of lactose and galactose by *Streptococcus mutans*: transport, toxicity, and carbon catabolite repression. J Bacteriol 192:2434–44.

30. Tong H, Zeng L, Burne RA. 2011. The EIIAB^Man^ PTS permease regulates carbohydrate catabolite repression in *Streptococcus gordonii*. Appl Environ Microbiol 77:1957–65.

31. Dong Y, Chen YY, Burne RA. 2004. Control of expression of the arginine deiminase operon of *Streptococcus gordonii* by CcpA and Flp. J Bacteriol 186:2511–4.

32. Loo CY, Corliss DA, Ganeshkumar N. 2000. *Streptococcus gordonii* biofilm formation: identification of genes that code for biofilm phenotypes. J Bacteriol 182:1374–82.

33. Culp DJ, Hull W, Schultz AC, Bryant AS, Lizarraga CA, Dupuis MR, Chakraborty B, Lee K, Burne RA. 2022. Testing of candidate probiotics to prevent dental caries induced by *Streptococcus mutans* in a mouse model. J Appl Microbiol 132:3853–3869.

34. Chen L, Chakraborty B, Zou J, Burne RA, Zeng L. 2019. Amino sugars modify antagonistic interactions between commensal oral streptococci and *Streptococcus mutans*. Appl Environ Microbiol 85:e00370–19.

35. Redanz S, Cheng X, Giacaman RA, Pfeifer CS, Merritt J, Kreth J. 2018. Live and let die: Hydrogen peroxide production by the commensal flora and its role in maintaining a symbiotic microbiome. Mol Oral Microbiol 33:337–352.

36. Mann AE, Chakraborty B, O’Connell LM, Nascimento MM, Burne RA, Richards VP. 2024. Heterogeneous lineage-specific arginine deiminase expression within dental microbiome species. Microbiology Spectrum 12:e01445–23.

37. Kreth J, Giacaman RA, Raghavan R, Merritt J. 2017. The road less traveled - defining molecular commensalism with *Streptococcus sanguinis*. Mol Oral Microbiol 32:181–196.

38. Jennings W, Epand RM. 2020. CDP-diacylglycerol, a critical intermediate in lipid metabolism. Chem Phys Lipids 230:104914.

39. Xu P, Alves JM, Kitten T, Brown A, Chen Z, Ozaki LS, Manque P, Ge X, Serrano MG, Puiu D, Hendricks S, Wang Y, Chaplin MD, Akan D, Paik S, Peterson DL, Macrina FL, Buck GA. 2007. Genome of the opportunistic pathogen *Streptococcus sanguinis*. J Bacteriol 189:3166–3175.

40. Jenkinson HF, Lala HC, Shepherd MG. 1990. Coaggregation of *Streptococcus sanguis* and other streptococci with *Candida albicans*. Infect Immun 58:1429–36.

41. Dutton LC, Paszkiewicz KH, Silverman RJ, Splatt PR, Shaw S, Nobbs AH, Lamont RJ, Jenkinson HF, Ramsdale M. 2015. Transcriptional landscape of trans-kingdom communication between *Candida albicans* and *Streptococcus gordonii*. Mol Oral Microbiol doi:10.1111/omi.12111.

42. Zeng L, Burne RA. 2010. Seryl-phosphorylated HPr regulates CcpA-independent carbon catabolite repression in conjunction with PTS permeases in *Streptococcus mutans*. Mol Microbiol 75:1145–58.

43. Fleming E, Lazinski DW, Camilli A. 2015. Carbon catabolite repression by seryl phosphorylated HPr is essential to *Streptococcus pneumoniae* in carbohydrate-rich environments. Mol Microbiol 97:360–80.

44. Deutscher J, Sauerwald H. 1986. Stimulation of dihydroxyacetone and glycerol kinase activity in Streptococcus faecalis by phosphoenolpyruvate-dependent phosphorylation catalyzed by enzyme I and HPr of the phosphotransferase system. J Bacteriol 166:829–36.

45. Zeng L, Burne RA. 2008. Multiple sugar: phosphotransferase system permeases participate in catabolite modification of gene expression in *Streptococcus mutans*. Mol Microbiol 70:197–208.

46. Abranches J, Nascimento MM, Zeng L, Browngardt CM, Wen ZT, Rivera MF, Burne RA. 2008. CcpA regulates central metabolism and virulence gene expression in *Streptococcus mutans*. J Bacteriol 190:2340–9.

47. Jurysta C, Bulur N, Oguzhan B, Satman I, Yilmaz TM, Malaisse WJ, Sener A. 2009. Salivary Glucose Concentration and Excretion in Normal and Diabetic Subjects. Journal of Biomedicine and Biotechnology 2009:430426.

48. Charrier V, Buckley E, Parsonage D, Galinier A, Darbon E, Jaquinod M, Forest E, Deutscher J, Claiborne A. 1997. Cloning and sequencing of two enterococcal *glpK* genes and regulation of the encoded glycerol kinases by phosphoenolpyruvate-dependent, phosphotransferase system-catalyzed phosphorylation of a single histidyl residue. J Biol Chem 272:14166–74.

49. Boulanger EF, Sabag-Daigle A, Thirugnanasambantham P, Gopalan V, Ahmer BMM. 2021. Sugar-Phosphate Toxicities. Microbiol Mol Biol Rev 85:e0012321.

50. Subedi KP, Kim I, Kim J, Min B, Park C. 2008. Role of GldA in dihydroxyacetone and methylglyoxal metabolism of Escherichia coli K12. FEMS Microbiol Lett 279:180–7.

51. Cornejo OE, Lefebure T, Pavinski Bitar PD, Lang P, Richards VP, Eilertson K, Do T, Beighton D, Zeng L, Ahn SJ, Burne RA, Siepel A, Bustamante CD, Stanhope MJ. 2013. Evolutionary and population genomics of the cavity causing bacteria *Streptococcus mutans*. Mol Biol Evol 30:881–93.

52. Saito M, Seki M, Iida K-i, Nakayama H, Yoshida S-i. 2007. A novel agar medium to detect hydrogen peroxide-producing bacteria based on the prussian blue-forming reaction. Microbiol Immunol 51:889–892.

53. Terleckyj B, Willett NP, Shockman GD. 1975. Growth of several cariogenic strains of oral streptococci in a chemically defined medium. Infect Immun 11:649–55.

54. Andrews S. 2015. FASTQC: A Quality Control tool for High Throughput Sequence Data. Babraham Institute.

55. Martin M. 2011. Cutadapt removes adapter sequences from high-throughput sequencing reads. EMBnetjournal doi:10.14806/ej.17.1.200.

56. Li H, Durbin R. 2009. Fast and accurate short read alignment with Burrows-Wheeler transform. Bioinformatics 25:1754–1760.

57. Quast C, Pruesse E, Yilmaz P, Gerken J, Schweer T, Yarza P, Peplies J, Glockner FO. 2013. The SILVA ribosomal RNA gene database project: improved data processing and web-based tools. Nucleic Acids Res 41:D590–6.

58. Li H, Handsaker B, Wysoker A, Fennell T, Ruan J, Homer N, Marth G, Abecasis G, Durbin R, Genome Project Data Processing S. 2009. The Sequence Alignment/Map format and SAMtools. Bioinformatics 25:2078–2079.

59. Quinlan AR, Hall IM. 2010. BEDTools: a flexible suite of utilities for comparing genomic features. Bioinformatics 26:841–2.

60. Franzosa EA, McIver LJ, Rahnavard G, Thompson LR, Schirmer M, Weingart G, Lipson KS, Knight R, Caporaso JG, Segata N, Huttenhower C. 2018. Species-level functional profiling of metagenomes and metatranscriptomes. Nat Methods 15:962–968.

61. Team RDC. 2010. R: A language and environment for statistical computing. R Foundation for Statistical Computing, Vienna, Austria.

62. Robinson MD, McCarthy DJ, Smyth GK. 2009. edgeR: A Bioconductor package for differential expression analysis of digital gene expression data. Bioinformatics doi:10.1093/bioinformatics/btp616.

