## Supplementary material for "Glycerol Metabolism Contributes to Competition by Oral Streptococci through Production of Hydrogen Peroxide": Includes Figures S1 to S5 and Tables S1 and S3.

### Supplemental Materials for “GLYCEROL METABOLISM CONTRIBUTES TO COMPETITION BY ORAL STREPTOCOCCI THROUGH PRODUCTION OF HYDROGEN PEROXIDE”

**Includes Figures S1 to S5 and Tables S1 and S3.**

**Fig. S1** Growth curves of oral streptococci and certain mutant derivatives incubated in FMC modified to contain various carbohydrate. Bacterial cultures prepared with BHI were diluted 100-fold into 200 μl FMC medium in a 96-well microtiter plate and covered with 60 μl mineral oil. Incubation of the plates was carried out in air (A to M, O) and the optical density ($OD_{600}$) of the cultures was monitored using a Synergy 2 (Biotek) plate reader maintained at 37°C. Fructose (Fru) and galactose (Gal) were each used at 20 mM, glycerol (Gly) at 5 mM or 40 mM, and catalase at 5 μg/ml. (N) Incubation in an anaerobic chamber abolished the low-level growth by *S. sanguinis* on glycerol observed in microaerophilic condition. Each strain was represented by at least three biological repeats, with their averages in $OD_{600}$ and standard deviations (error bars) being presented here.

A

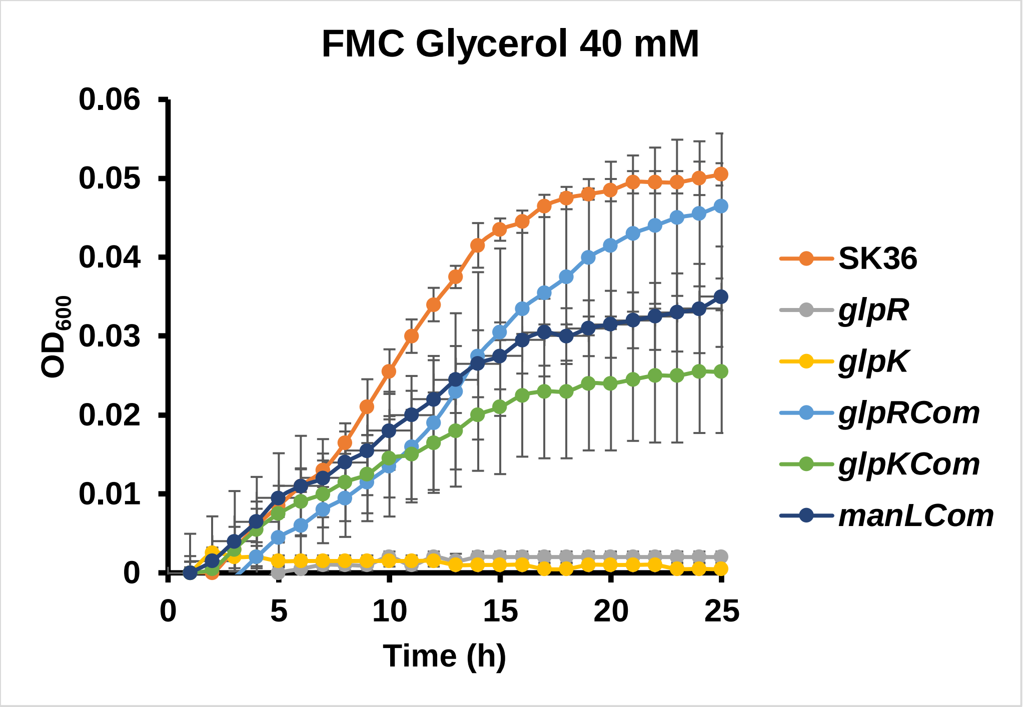

B

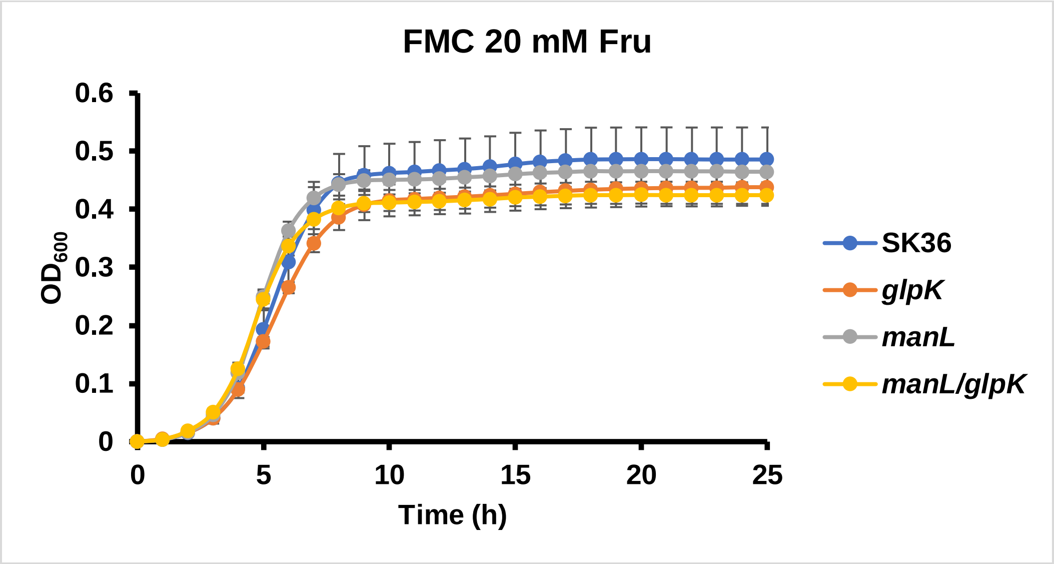
FMC 20 mM Fru
0.6
0.5
0.4
0.3
0.2
0.1
0
$OD_{600}$
0
5
10
15
20
25
Time (h)
SK36
glpK
manL
manL/glpK

C

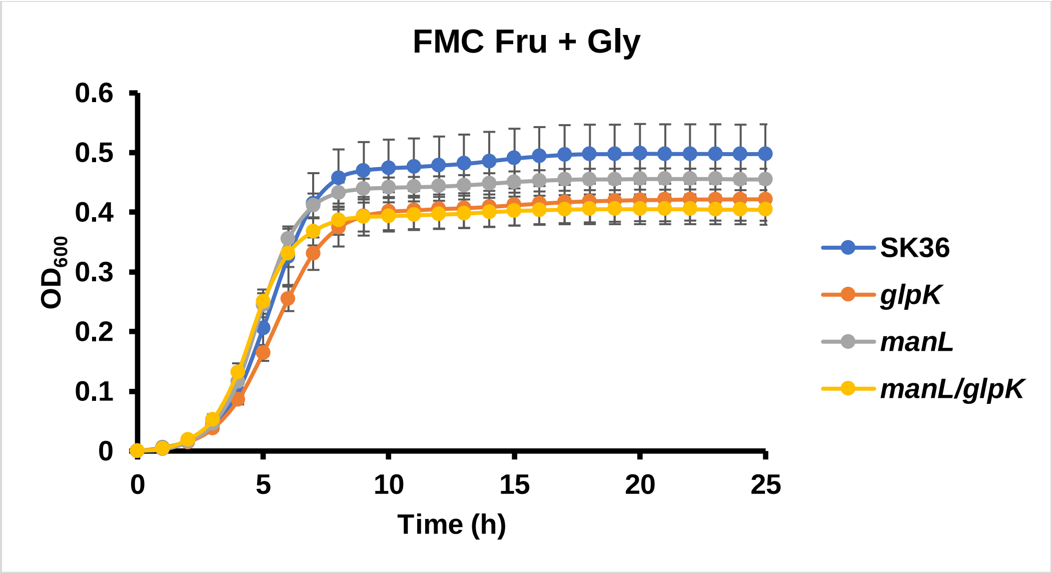
FMC Fru + Gly
0.6
0.5
0.4
0.3
0.2
0.1
0
$OD_{600}$
0
5
10
15
20
25
Time (h)
SK36
glpK
manL
manL/glpK

D

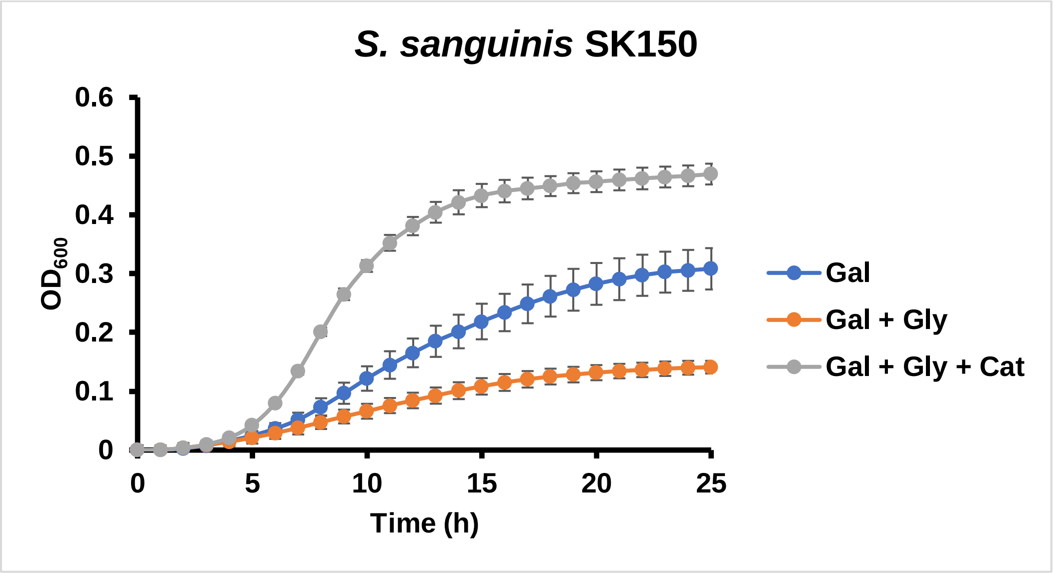
S. sanguinis SK150
0.6
0.5
0.4
0.3
0.2
0.1
0
$OD_{600}$
0
5
10
15
20
25
Time (h)
Gal
Gal + Gly
Gal + Gly + Cat

E

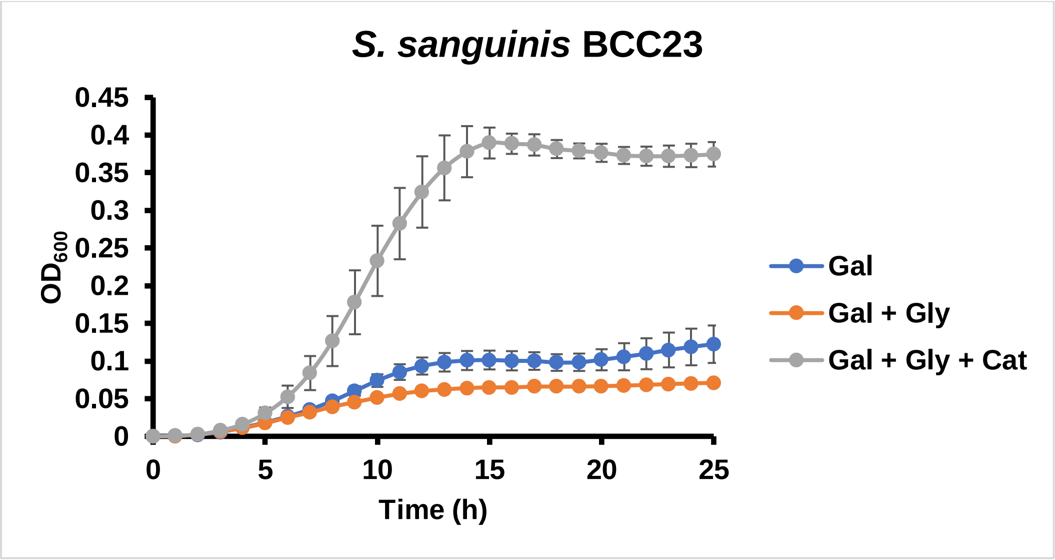
S. sanguinis BCC23
0.45
0.4
0.35
0.3
0.25
0.2
0.15
0.1
0.05
0
$OD_{600}$
0
5
10
15
20
25
Time (h)
Gal
Gal + Gly
Gal + Gly + Cat

F

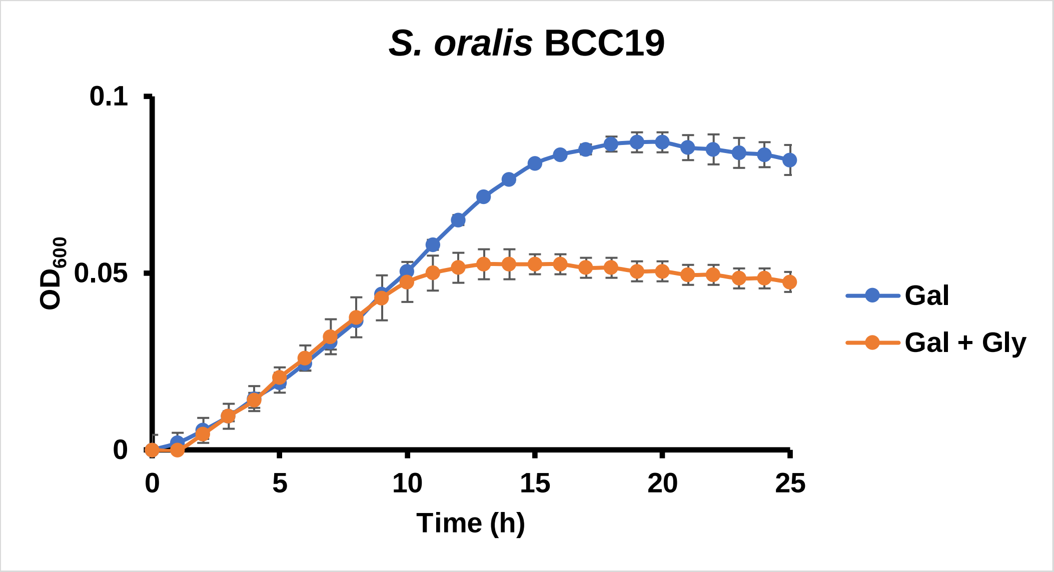
S. oralis BCC19
0.1
0.05
0
OD600
0
5
10
15
20
25
Time (h)
Gal
Gal + Gly

G

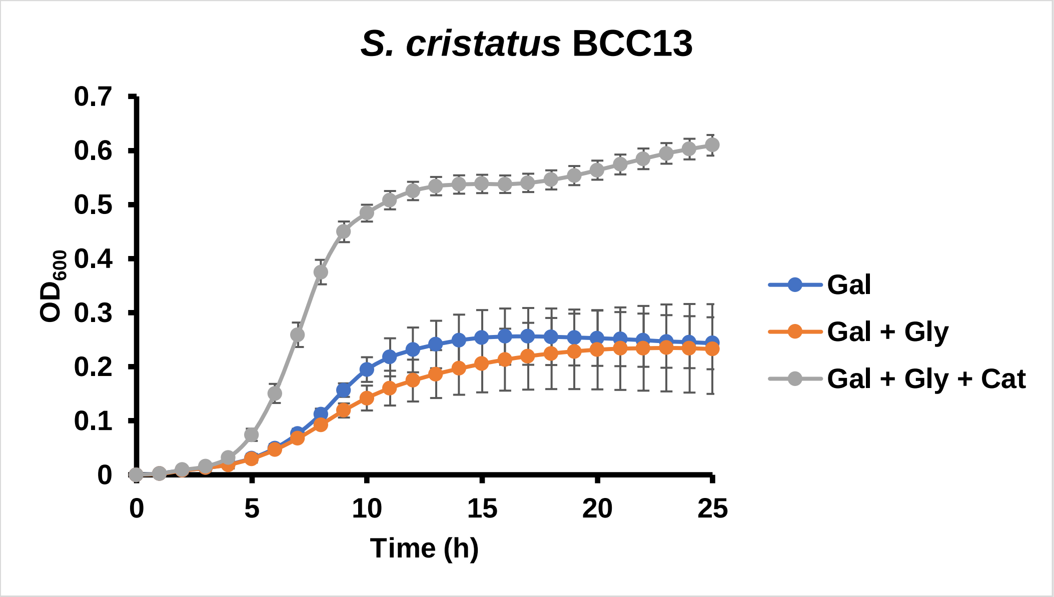
S. cristatus BCC13
0.7
0.6
0.5
0.4
0.3
0.2
0.1
0
OD600
0
5
10
15
20
25
Time (h)
Gal
Gal + Gly
Gal + Gly + Cat

H

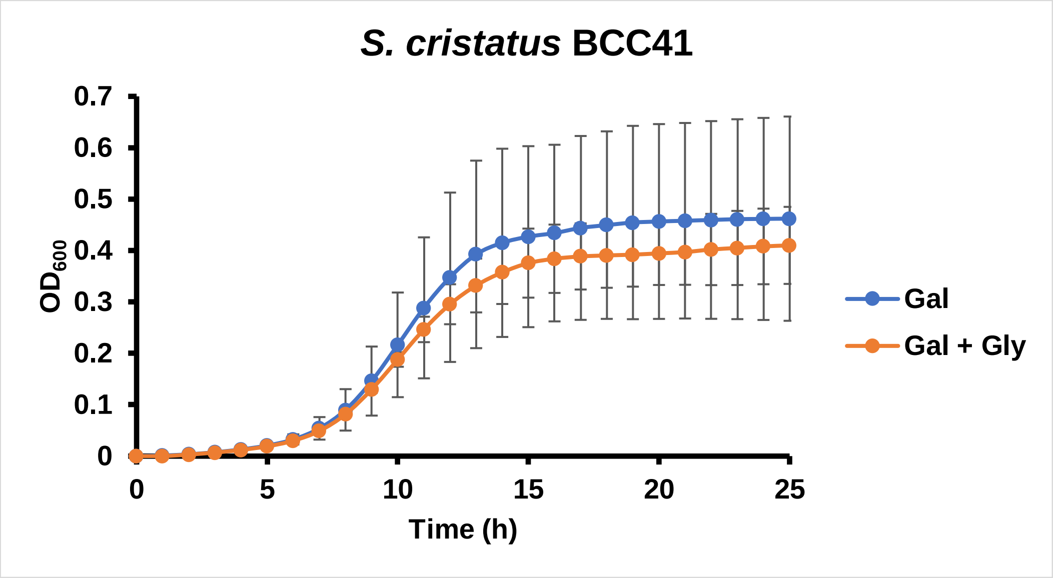

S. cristatus BCC41
0.7
0.6
0.5
0.4
0.3
0.2
0.1
0
OD600
0
5
10
15
20
25
Time (h)
Gal
Gal + Gly


I

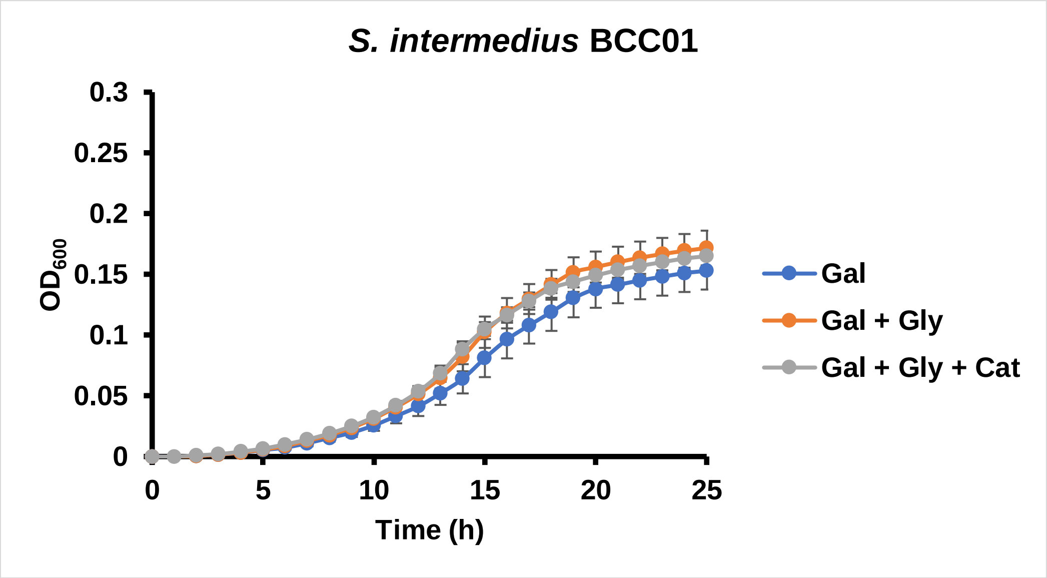

S. intermedius BCC01
0.3
0.25
0.2
0.15
0.1
0.05
0
OD600
0
5
10
15
20
25
Time (h)
Gal
Gal + Gly
Gal + Gly + Cat

J

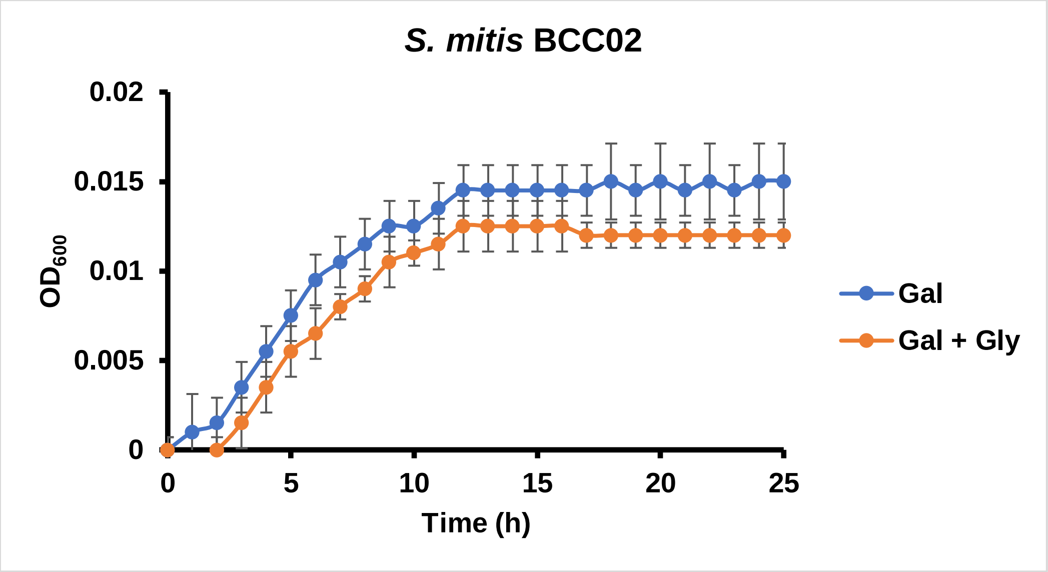
S. mitis BCC02
0.02
0.015
0.01
0.005
0
OD600
0
5
10
15
20
25
Time (h)
Gal
Gal + Gly

K

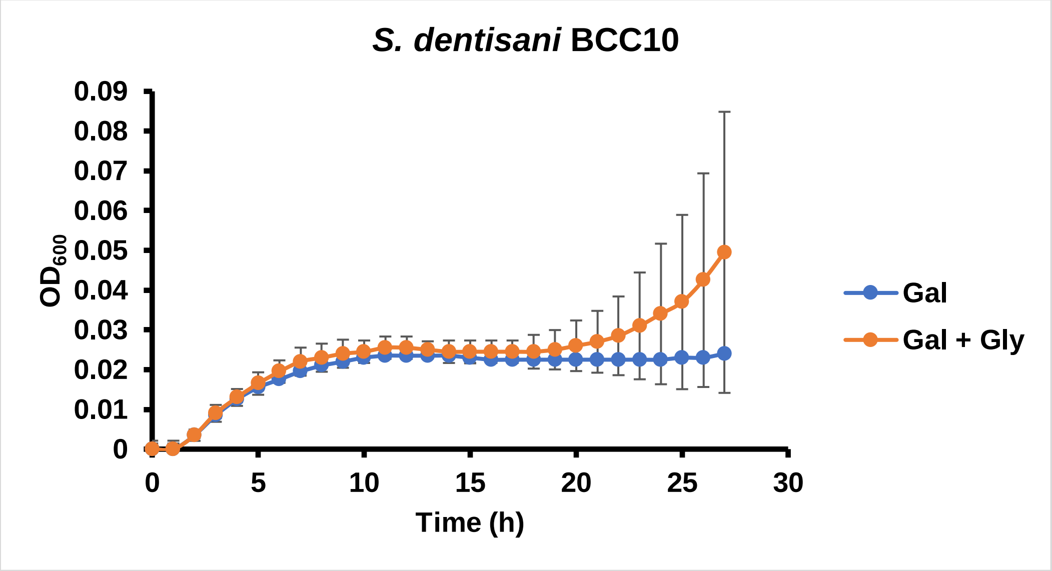
S. dentisani BCC10
0.09
0.08
0.07
0.06
0.05
0.04
0.03
0.02
0.01
0
OD600
0
5
10
15
20
25
30
Time (h)
Gal
Gal + Gly

L

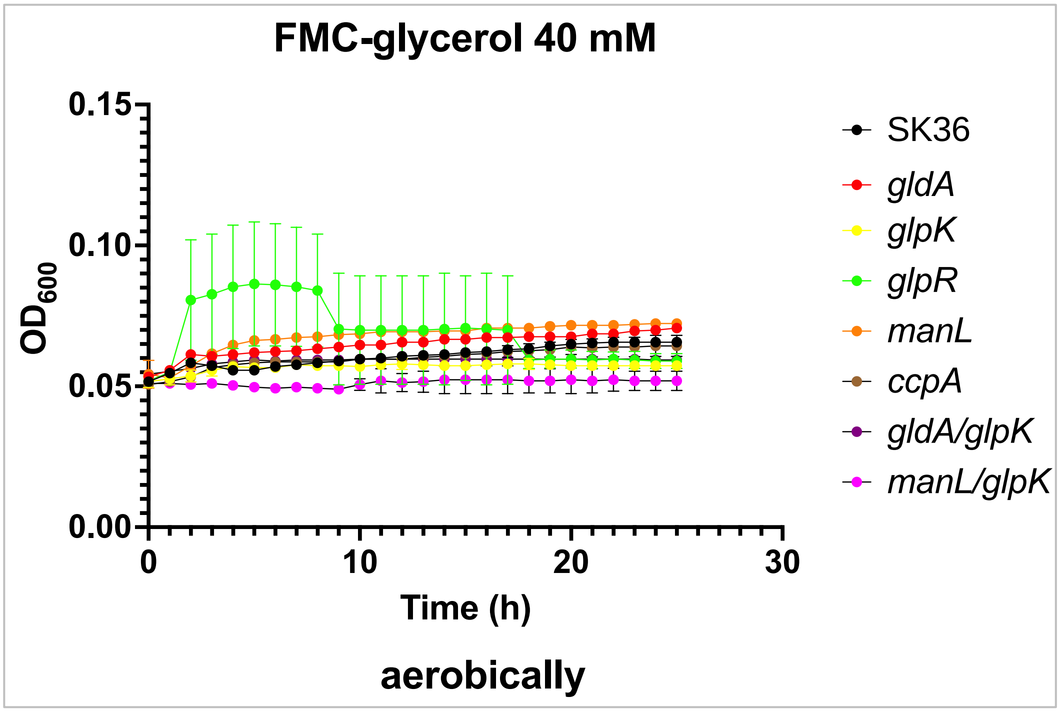
FMC-glycerol 40 mM
0.15
0.10
0.05
0.00
OD600
0
10
20
30
Time (h)
SK36
gldA
glpK
glpR
manL
ccpA
gldA/glpK
manL/glpK
aerobically

M

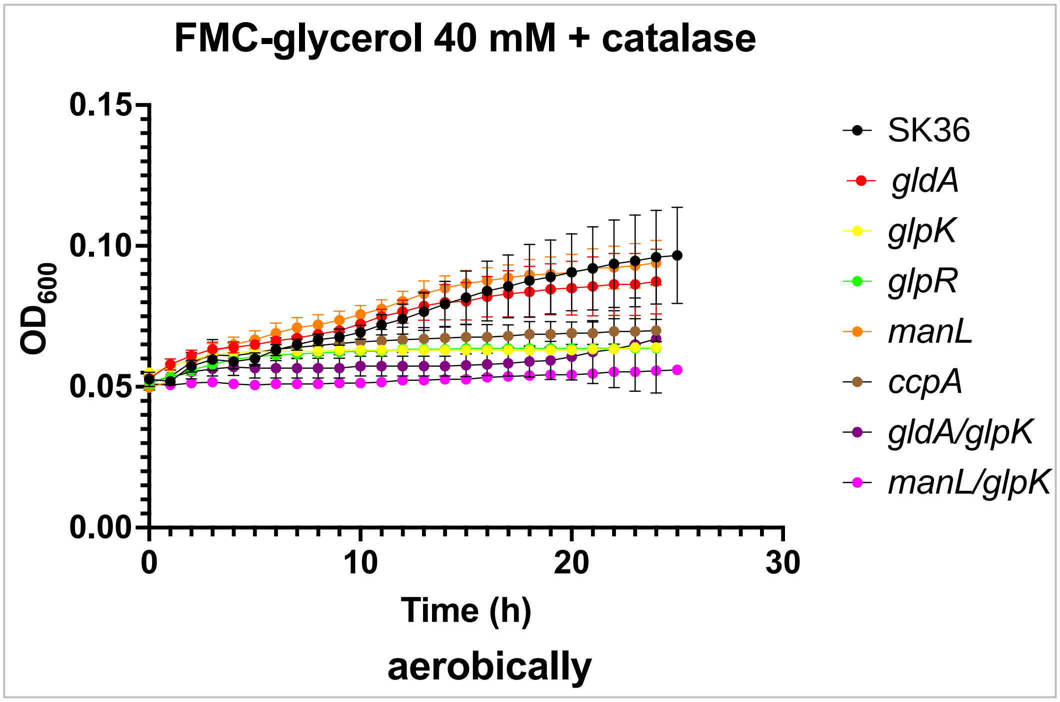
FMC-glycerol 40 mM + catalase
0.15
0.10
0.05
0.00
OD600
0
10
20
30
Time (h)
aerobically
SK36
gldA
glpK
glpR
manL
ccpA
gldA/glpK
manL/glpK

N

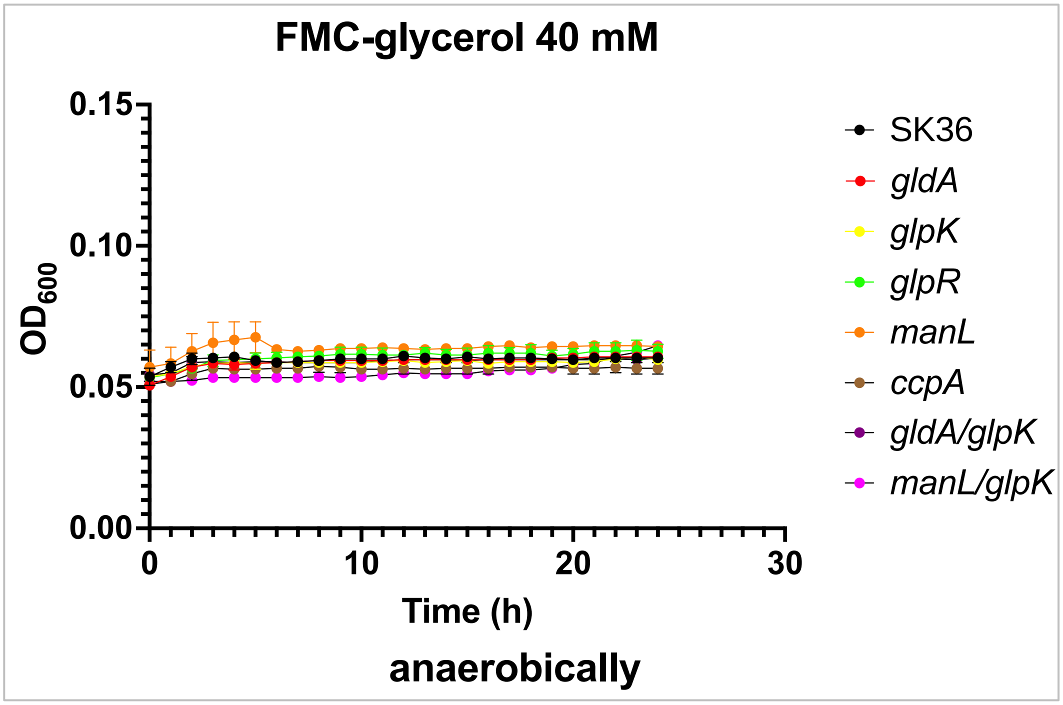
FMC-glycerol 40 mM
0.15
0.10
0.05
0.00
OD600
0
10
20
30
Time (h)
anaerobically
SK36
gldA
glpK
glpR
manL
ccpA
gldA/glpK
manL/glpK

O

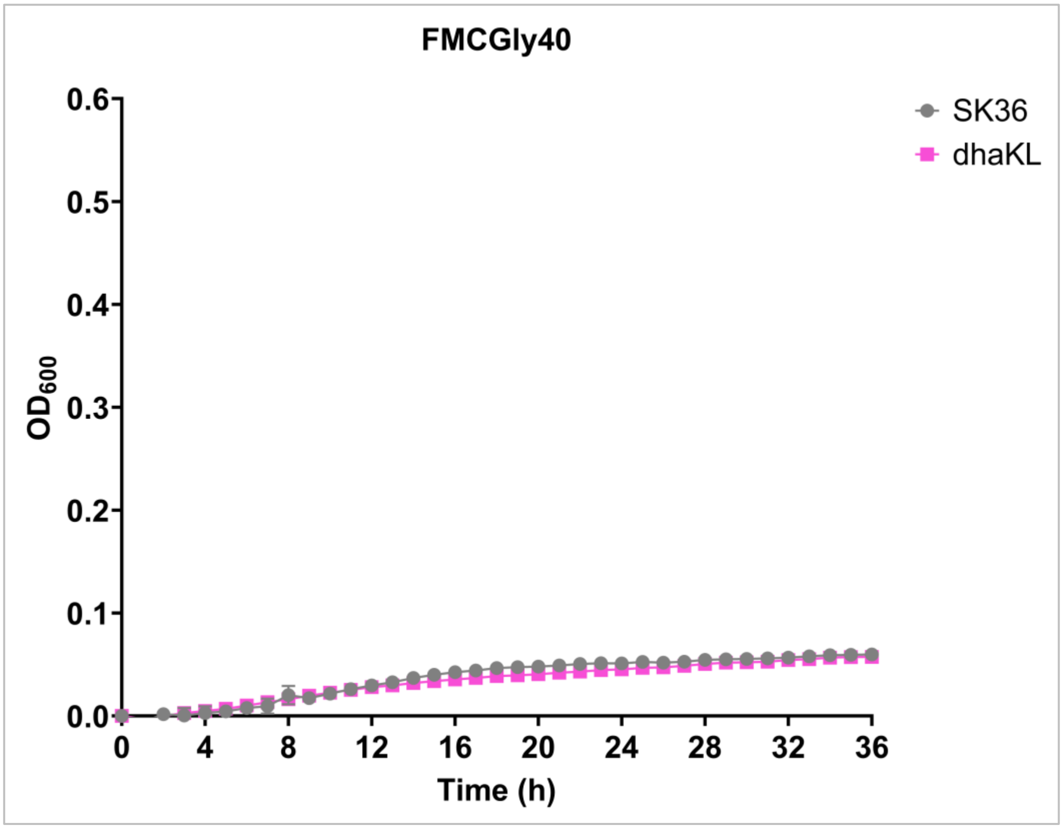
FMCGly40
0.6
0.5
0.4
0.3
0.2
0.1
0.0
OD600
0
4
8
12
16
20
24
28
32
36
Time (h)
SK36
dhaKL

**Fig. S2 Prussian Blue assay for detection of $H_2O_2$.** Bacterial cultures were placed on surface of Prussian Blue agar plates containing TY medium supplemented with 20 mM of glucose or glycerol, incubated overnight in an ambient-air incubator supplemented with 5% $CO_2$. Strains included (A) the WT SK36 and Δ*dhaKL*, and (B) mutants deficient in *spxB*, *manL* (1918), *glpK* (1826), *glpO* (1827), and *glpF* (1828) that were constructed previously (1).

A

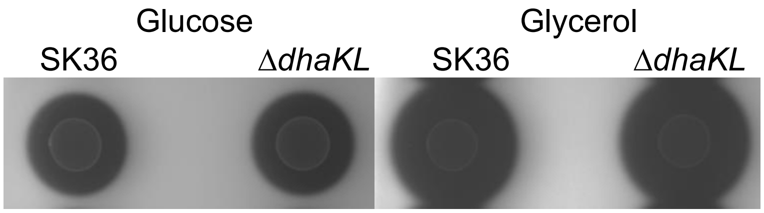


B

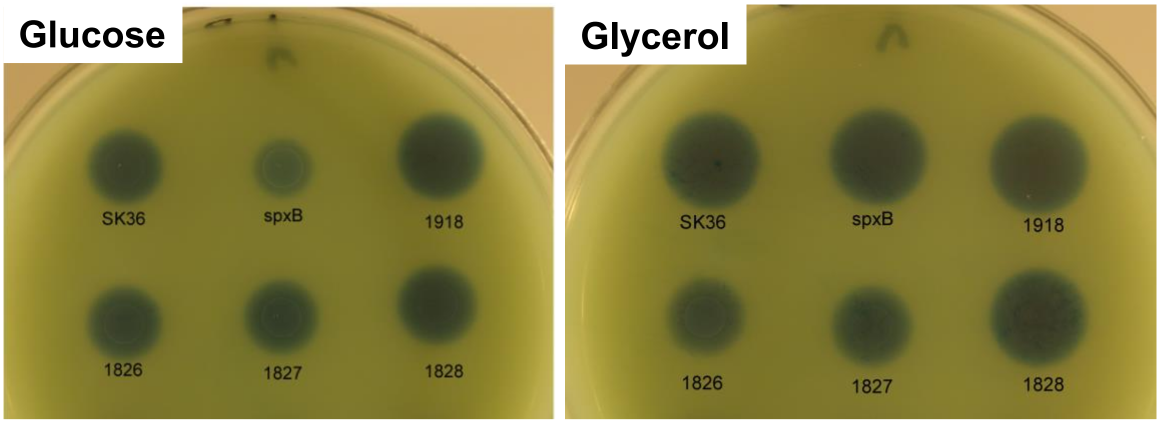

**Fig. S3** Measurements of glycerol in various media used in this study and spent media from cultures of *Candida albicans. C. albicans* strain SC5314 was inoculated in BHI for overnight growth. The yeast cells were washed twice with FMC base medium and diluted 20-fold into FMC supplemented with 20 mM of glucose (Glc), galactose, N-acetylglucosamine (GlcNAc), or mannose. The cultures were kept at 37°C with shaking at 200 RPM till $OD_{600}$ reached 1 or stopped increasing. The culture supernatant was then harvested by centrifugation, filtered and frozen before assay. Each sample was measured 4 times using a glycerol assay kit from Neogen (Lansing, MI).

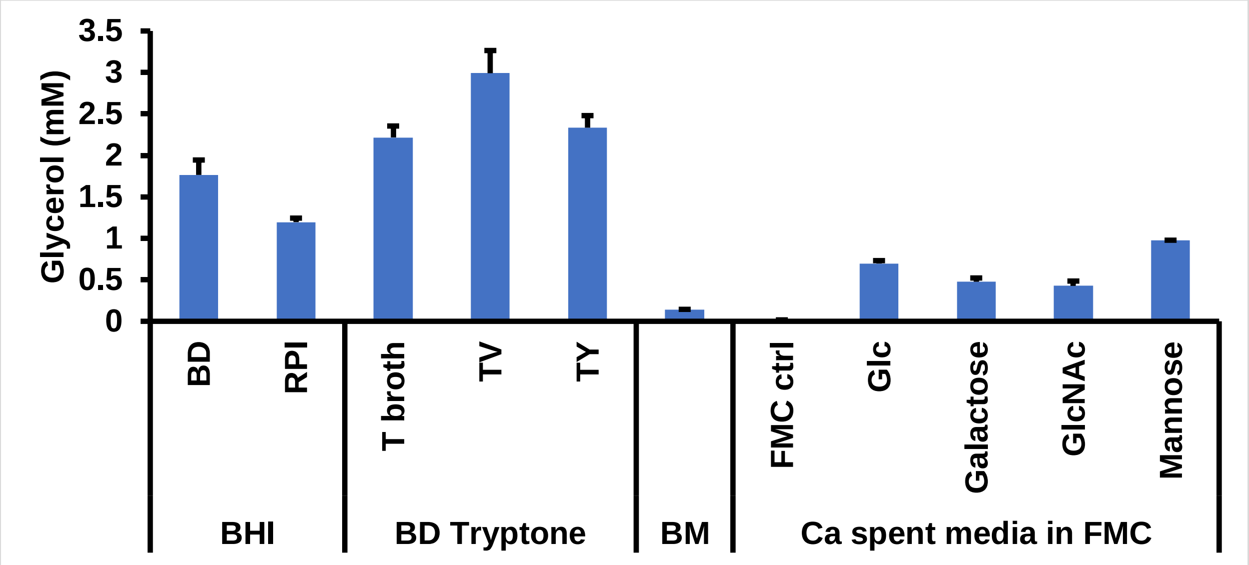

**Fig. S4** Additional competition assays between *S. sanguinis* and *S. mutans* in a two-species biofilm model. SK36 or its isogenic mutant Δ*glpK* was mixed with an equal volume of *S. mutans* UA159 culture and inoculated, at 1:100 ratio, into a BMGS medium held in a 24-well plate with a hydroxyapatite disc kept in an (A) aerobic and (B) anaerobic atmosphere. After 24 h of incubation, the culture supernatant was replaced with BM base, BM containing 18 mM glucose (Glc), or BM with 36 mM glycerol (Gly). After another day of incubation, the biofilm was harvested for CFU enumeration. Each strain was represented by 4 separate cultures and each condition 4 biofilm samples. The average CFU and standard deviation (error bars) of each species was used to plot the graph and for statistics (One-way ANOVA followed by Tukey's multiple comparisons; *, $P$ <0.05; ****, $P$ <0.0001).

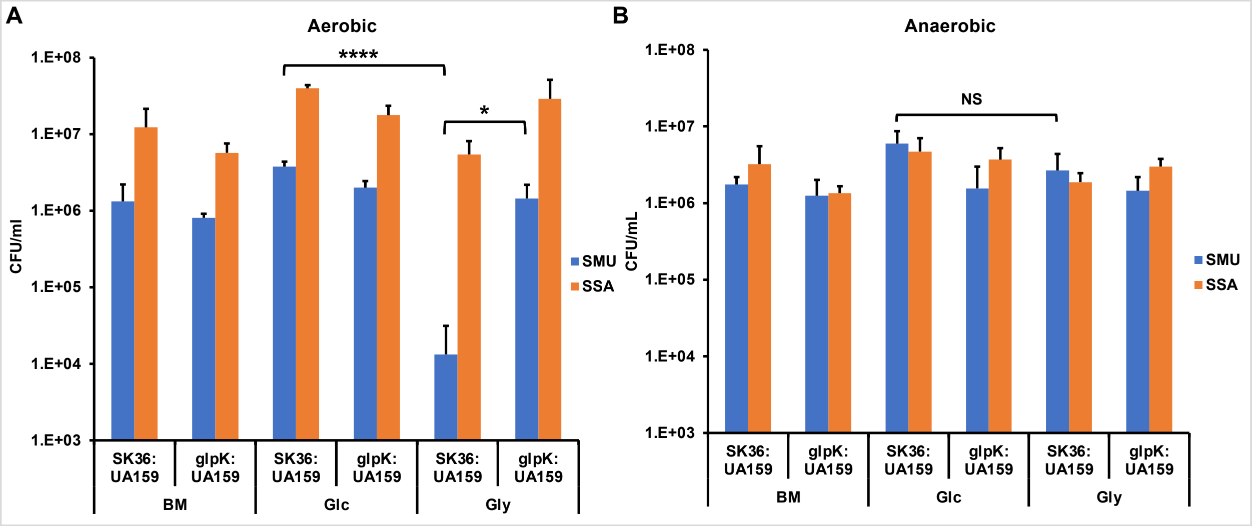

**Fig. S5** Identification of CcpA-binding *cre* element (in blue) upstream of *glpK* and *dhaK*.

attcatcaggcccagcaggttcctatctattatttaaaaaatgacttggatttggaagat
ttggctcaaatccgtcacatgatatttcattaagaaacttggagaaatccaggtttcttt
gtgattttcgcacaatctcctcagatttttttaaaatttaaaaaaagttgataaaaa
agaaagcgttttattttatatactagttactgtagaggaaacatcctcaagatttttactagg
aggacagttcATGTCACAAGAAAAATACATCATGGCCATCGACCAAGGAACTACTAGTTC (SSA_1826/*glpK*)

tgttctttcagctgggcagtattgcgtttcatgtatctcctccaaaagtatacactgtct
acttatctatctttagtatacactgtatactttttattgtcaaataatttgatgctttat
tctagtctaagatgtggcagaatttgctttcggtaagataaaatgttaaaataatatcca
aatgatagcgctttcttttttttggggtggattgctcgttgtaaagaaacgctgagaaacg
gaggacacctATGAAGAAAATTATCAACGACCCAACAGCGGTTGTGGACGAGATGCTGGA (SSA_0049/*dhaK*)

**Table S1** Relevant genes showing differential expression in the Δ*manL* background as indicated by RNA-seq (2).

| Name | Number | Annotation | Δ*manL*/SK36 | FDR |
|---|---|---|---|---|
| *glpK* | SSA_1826 | glycerol kinase GlpK | 90.7 | 0 |
| *glpO* | SSA_1827 | type 1 glycerol-3-phosphate oxidase | 88.5 | 0 |
| *glpF* | SSA_1828 | aquaporin family protein | 71.4 | 0 |
| *dhaK* | SSA_0049 | dihydroxyacetone kinase subunit 1 | 9.9 | 8.3e-140 |
| *dhaL* | SSA_0050 | dihydroxyacetone kinase subunit L | 12.0 | 8.2e-123 |
| *dhaM* | SSA_0051 | PTS-dependent dihydroxyacetone kinase phosphotransferase subunit | 12.7 | 9.1e-113 |
| *gldA* | SSA_0287 | glycerol dehydrogenase | 0.5 | 2.0e-9 |

**Table S3** Primers used in this study.

| Primers | Sequence | Purpose |
|---|---|---|
| SSA_glpK-1 | GGA CAG CGA TTG GAC TTG G | Δ*glpK* |
| SSA_glpK-2GA | GCC ATT TAT TAT TTC CTCT CTT TTA C TTG TGA CAT GAA CTG TCC TCC | Δ*glpK* |
| SSA_glpK-3GA | ATA TTT TAC TGG ATG AAT TGT TTT AGT AGA CGA CGA CTG ATA CTG ATT TAT | Δ*glpK* |
| SSA_glpK-4 | CCA GGT CAC CAG TGT AGT C | Δ*glpK* |
| SSA_glpKCom-2GA | GCC ATT TAT TAT TTC CTT CCT CTT TTA CAG TAT CAG TCG TCG ATT TCT GCA | Δ*glpK*Com |
| SSA_glpR-1 | CAT GAC CTT GGG TGG CGA AT | Δ*glpR* |
| SSA_glpR-2GA | GCC ATT TAT TAT TTC CTT CCT CTT TTA GCT GCT GTA AAT AAG ACA GGA CA | Δ*glpR* |
| SSA_glpR-3GA | ATA TTT TAC TGG ATG AAT TGT TTT AGT AGA GGC TCA AAT CCG TCA CAT GA | Δ*glpR* |
| SSA_glpR-4 | CCA TAC CAG GCT CAA AAG CTA ACT | Δ*glpR* |
| SSA_glpRCom-2GA | GCC ATT TAT TAT TTC CTT CCT CTT TTA GTT TCT TAA TGA AAT ATC ATG TGA CGG A | Δ*glpR*Com |
| SSA_glpF-1 | GCG TCC TCT AAT CGC AGG A | Δ*glpF* |
| SSA_glpF-4 | GAG ACA GTG GCG GTC AAT T | Δ*glpF* |
| SSA_glpF-2GA | GCC ATT TAT TAT TTC CTT CCT CTT TTA GAG CAA GAG CAG TGT TCC T | Δ*glpF* |
| SSA_glpF-3GA | ATA TTT TAC TGG ATG AAT TGT TTT AGT AGA GCA GTC CTA GTC TTC GGT T | Δ*glpF* |
| SSA_gldA-1 | CCT CAG ACC CTG TCT AAG GA | Δ*gldA* |
| SSA_gldA-2GA | GCC ATT TAT TAT TTC CTT CCT CTT TTA CGA GAC GGG CTT GCA AAA ATT | Δ*gldA* |
| SSA_gldA-3GA | ATA TTT TAC TGG ATG AAT TGT TTT AGT AGA CCA TCC TAG CGG TTG ACC AA | Δ*gldA* |
| SSA_gldA-4 | GGC TTG TCC TTG TCC GTA CCT | Δ*gldA* |
| SSA_dhaK-1 | GTC TTG GCC GAT AGG ACT GT | Δ*dhaKL* |
| SSA_dhaK-2GA | GCC ATT TAT TAT TTC CTT CCT CTT TTA CCG CTG TTG GGT CGT T | Δ*dhaKL* |
| SSA_dhaL-3GA | ATA TTT TAC TGG ATG AAT TGT TTT AGT AGA GAG GCA GGC TGA TGG CA | Δ*dhaKL* |
| SSA_dhaL-4 | CGC GGT GAC TTC TTC AAG A | Δ*dhaKL* |
| SSA_spxB-1 | GGAGCAGGCAAAATTGCA | Δ*spxB* |
| SSA_spxB-2GA | GCCATTTATTATTTCCTTCCTCTTTTACGTTGAGCATTGC TGCAGAT | Δ*spxB* |
| SSA_spxB-3GA | ATATTTTACTGGATGAATTGTTTTAGTAGACTTGGAAGAA GAAGGATTGCAATCA | Δ*spxB* |
| SSA_spxB-4 | ACAATCTCATTATTCCTATCTTCTATCTGTT | Δ*spxB* |
| SSA_gldA-AS | TTAAAAGCGACATGCAGCAC | RT-qPCR |
| SSA_glpK-S | GAT ACC TGG CTG GTT TGG AA | RT-qPCR |
| SSA_glpK-AS | CAA AAT CTC GTC GTC CCA TT | RT-qPCR |
| SSA_gldA-S | GCCATCCAGTCTTGCTTTGT | RT-qPCR |

| | | |
|---|---|---|
| SSA_dhaL-S | TTC AAG CTG TTT CCA TGC AG | RT-qPCR |
| SSA_dhaL-AS | TTG CCG TCT TTT TCA GCT TT | RT-qPCR |
| SSA_spxB-AS | TCT TCC AAG AAG AGG CGG AAT GGT | RT-qPCR |
| SSA_spxB-S | ATC ACT CAA CAC CGT CCA CTT CCA | RT-qPCR |

#### Reference


1. Xu P, Ge X, Chen L, Wang X, Dou Y, Xu JZ, Patel JR, Stone V, Trinh M, Evans K, Kitten T, Bonchev D, Buck GA. 2011. Genome-wide essential gene identification in *Streptococcus sanguinis*. Sci Rep 1:125.
2. Zeng L, Walker AR, Burne RA, Taylor ZA. 2022. Glucose phosphotransferase system modulates pyruvate metabolism, bacterial fitness, and microbial ecology in oral streptococci. J Bacteriol 205:e0035222.
